# A non-invasive approach to awake mouse fMRI compatible with multi-modal techniques

**DOI:** 10.1101/2025.01.23.634586

**Authors:** Sam Laxer, Amr Eed, Miranda Bellyou, Peter Zeman, Kyle M Gilbert, Mohammad Naderi, Ravi S Menon

**Affiliations:** Centre for Functional and Metabolic Mapping, Western University, London, Ontario, Canada; Department of Medical Biophysics, Western University, London, Ontario, Canada

**Keywords:** fMRI, awake mouse imaging, non-invasive restraint

## Abstract

Mouse functional magnetic resonance imaging (fMRI) studies contribute significantly to basic fundamental and translational neuroscience research. Performing fMRI in awake mice could facilitate complex tasks in the magnet and improve translational validity by avoiding anesthesia-related neural and neurovascular changes. Existing surgical approaches provide excellent motion control but are not desirable for all experiments aiming to scan awake mice. These include studies with transgenic mouse lines that are vulnerable to anesthesia or mice in longitudinal studies involving cognition. To address these needs, we propose a non-invasive restraint to scan mice in the awake state. The restraint was designed to be compatible with brain stimulation and recording approaches often combined with fMRI. It was evaluated on the basis of motion, fMRI data quality, and animal stress levels, and compared to a conventional headpost restraint. We found the proposed approach was effective at restraining mice across a broad range of weights without the need for any anesthesia for setup. The non-invasive restraint led to higher data attrition after censoring high motion volumes, but by acquiring roughly 25% more data we could obtain comparable network spatial specificity to the headpost approach. Our results demonstrate a simple open-source head restraint that can be used for awake mouse fMRI for certain cohorts, and we establish suitable acclimation and scanning protocols for use with this restraint.

## 1 Introduction

Functional magnetic resonance imaging (fMRI) is an important tool in neuroscience for characterizing brain-wide networks in healthy animals and disease models. Mice have been particularly informative due to the wide range of available transgenic lines thought to have translational value in studies of human disease (Mandino et al., 2020, 2021; N. Xu et al., 2022). However, motion needs to be minimized to acquire useful data (Power et al., 2012). To control motion, there are currently 3 main approaches: 1) using anesthesia, 2) scanning awake using a surgically implanted headpost/head plate, and 3) scanning awake using a non-invasive restraint.

The use of anesthesia to immobilize mice has been common practice in structural and functional MRI research for decades. However, anesthesia can be problematic in certain scenarios. An increasing number of fMRI studies have shown its confounding effects on neurovascular coupling and network architecture (Desai et al., 2011; Dinh et al., 2021; Gao et al., 2017; Gutierrez-Barragan et al., 2022; Z. Han et al., 2019; Tsurugizawa & Yoshimaru, 2021). These effects are also observed in fMRI studies of nonhuman primates (Hori et al., 2020) and humans (Luppi et al., 2025). These effects depend on the variability of each animal’s response to anesthetic type, depth and duration which leads to network variability in and between animal studies and has implications for the translational validity of fMRI studies performed under anesthesia.

The vast majority of fMRI studies in anesthetized mice have been conducted in wildtype animals which are generally robust, but many transgenic lines especially those that model ageing, neurodevelopmental or neurodegenerative disorders, have higher mortality under anesthesia both during and after anesthesia (Schuetze et al., 2019). Additionally, there is a large literature on cognitive deficits in humans after one or more episodes of general anesthesia, and durable cognitive deficits are also becoming apparent in animal models (Guo et al., 2023). While the mechanisms for this are unclear, and the literature is equivocal, it appears that injectable anesthetics have better cognitive outcomes (Mardini et al., 2017) than inhalation anesthetics. Inhalation anesthetics appear to accelerate the accumulation of neuropathology or increased neurotoxicity in transgenic models of neurodegeneration (Bianchi et al., 2008; Feng et al., 2016; F. Han et al., 2021; He et al., 2024; Miao et al., 2018; Tang & Eckenhoff, 2013) which could be an additional causative factor for the cognitive declines observed. Adding to the difficulties in interpreting results, these effects are also sex-dependent in either a detrimental manner (Zhang et al., 2017) or, in the case of halothane, with differential protection/detriment in different brain areas for females (Tang et al., 2011). In general, the literature suggests that anesthesia should be avoided in these models.

Much of the research in our lab centers on the longitudinal scanning of newly developed humanized transgenic mouse models of neurodegenerative disorders such as Alzheimer’s and synucleinopathies. Such studies involve repeated structural and functional imaging of animals at multiple time points over a period of 16-20 months, often using implanted cannulas for optogenetics or fibre photometry. These animals are also evaluated at the same time points using sophisticated touchscreen cognitive tests that parallel those used in humans (Dumont et al., 2025). These conditions pose certain challenges as outlined earlier. To avoid the potential for cognitive decline from repeated anesthesia, and the reported enhancement of the disease pathologies we are trying to investigate, we examined various approaches for longitudinal scanning of these vulnerable cohorts awake.

One effective strategy involves utilizing surgically implanted headposts or head plates as a fixation point on the skull (Chen et al., 2020; Gutierrez-Barragan et al., 2022). This approach avoids the negative effects of repeated anesthesia (after the initial surgery done under anesthesia however) and in some cases allows for comparable motion control to anesthesia. Despite these benefits, the use of a headpost poses challenges in our application. Ideally, we wish to avoid operating on vulnerable transgenic mouse lines to implant a headpost (Szilagyi et al., 2018; Wu et al., 2020). There is evidence that even a single exposure to anesthesia can cause impairment (Zhou et al., 2024). One solution is to develop non-invasive awake imaging approaches that forego the use of a headpost or head plate (and the associated surgery).

Non-invasive mouse restraints have been developed in the past decade (Fadel et al., 2022; Ferris, 2022; Harris et al., 2015; Madularu et al., 2017). Amongst the current designs, most seem to suffer from cumbersome animal setup procedures, often requiring brief administration of anesthesia to position subjects in the MRI holder. Further, some of these designs are not compatible with the small inner diameter size of high-performance gradient inserts found in many rodent MRI systems (Madularu et al., 2017). Thus, there remains a need for a simple, effective non-invasive restraint ideally requiring no anesthesia for animal setup.

A major consideration with awake imaging that is not encountered when using anesthesia is the stress encountered by the animal in the restraint and the stress associated with being within the hostile environment of the MRI scanner. Animal stress has been shown to impact neurovascular responses and produce stress-related motion, while also imposing a negative impact on the animal’s wellbeing (Low et al., 2016). Consequently, a large effort has been made to develop restraint features and associated procedures to minimize stress. To this end, various labs have developed acclimation procedures to gradually expose subjects to restraint and the scanning environment (i.e., dark and loud environment simulating the auditory and visual experience of the scanner) (Chen et al., 2020; Desjardins et al., 2019; Dinh et al., 2021; Gutierrez-Barragan et al., 2022; Takata et al., 2018). Additionally, care has been taken to design restraints such that subject comfort is improved to further reduce stress (W. Xu et al., 2022).

Here, we present a novel approach to non-invasive awake-mouse MRI with a focus on compatibility with multi-modal experiments – all while remaining small enough to fit inside the 60-mm-diameter gradient coil found in many small-bore animal MRI scanners. The developed restraint allows for effective motion control and no need for anesthesia during animal setup. Motion and dual-regression-based resting state network robustness were evaluated in comparison with a gold-standard headpost restraint approach. Lastly, stress was analyzed by measuring weight and corticosterone (CORT) changes between three different acclimation procedures in both sexes.

## 2 Methods

### 2.1 Ethics statement

All experimental methods described were performed in accordance with the guidelines of the Canadian Council on Animal Care policy on the care and use of experimental animals and an ethics protocol approved by the Animal Care Committee of The University of Western Ontario.

### 2.2 Subjects and groups

This study involved 40 wildtype C57BL/6 mice from Jackson Laboratories (n = 13) and Charles River Laboratories (n = 27). Mice were divided into 4 groups (Table 1) with each group having roughly equally proportions of males and females. Groups 1, 2 and 3 underwent the non-invasive imaging protocol (with differing acclimation lengths) and Group 4 underwent a conventional headpost imaging protocol with the same acclimation protocol as the non-invasive 9-day protocol as shown in Table 2. Animals in Groups 1, 2, and 3 were single-housed in a facility with a 12-hour light/dark cycle (light from 7:00 am – 7:00 pm) and fed food and water ad-libitum. Animals were single-housed to replicate the treatment for cannula-implanted mice who are single-housed to minimize the risk of mouse cage mates damaging/removing each other’s cannulas. Mice in Group 4 (acquired previously) were group-housed. Importantly, mice of varying age and weight ranges were intentionally selected to validate the restraint on a diverse set of mice. Young mice tend to be smaller and lighter, with older mice being larger and heavier.

**Table 1.** Subject information. The first 4 rows display the sample sizes in each group (m = males, f = females). The wide age/weight range was to validate the restraint on mice of different sizes. Age and weight ranges were found by taking the lightest and heaviest mouse from the initial handling for acclimation to the last scanning day. The uncertainty of ± 1 week accounts for mice arriving at the facility from 6-8 weeks old.

| Parameter | Value |
| --- | --- |
| Group 1 (Non-invasive – 4-day protocol) | 4 m + 5 f |
| Group 2 (Non-invasive – 9-day protocol) | 4 m + 5 f |
| Group 3 (Non-invasive – 13-day protocol) | 7 m + 8 f |
| Group 4 (Headpost – 9-day protocol) | 4 m + 3 f |
| Strain | C57BL/6 |
| Age range | $8 \pm 1$ to $55 \pm 1$ weeks |
| Weight range | 16 – 37 g |

### 2.3 Restraint designs

The non-invasive restraint is almost entirely 3D printed of polycarbonate (Figure *1*). It consists of a cylindrical body tube, a neck plate to keep the subject in position, and the combination of a bite bar and two head arches (one anterior at the snout and one posterior at the caudal end of the skull) to stabilize the head in all three dimensions. An in-house built 2 cm single-loop transmit/receive coil was designed to fit over top of the head after the arches were secured.

**Figure 1.**
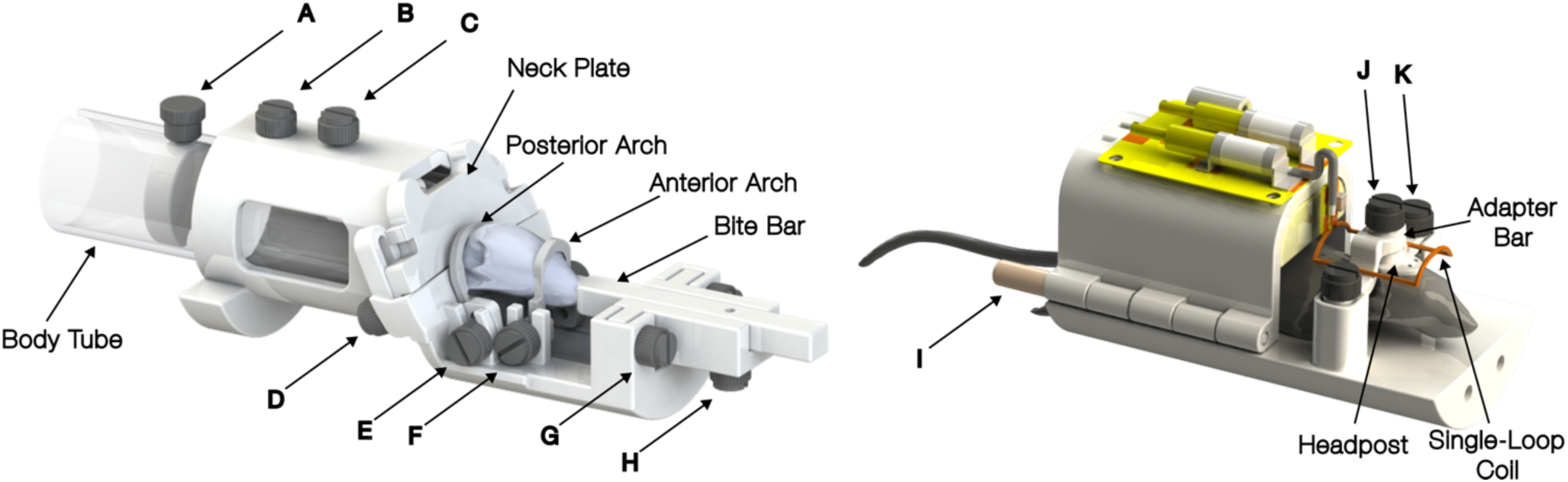
CAD models of the non-invasive restraint (left) and headpost restraint (right). The single loop coil used for single transmit/receive is not shown on the non-invasive restraint to allow clear visualization of the arches over the mouse head. A) Screw to position tail plate within body tube. B) Screw to secure body tube in restraint. C) Screw to hold coil (not shown). D) Screw to tighten neck plate (along with additional screw on other side). E) Screw to tighten posterior arch around posterior head (along with additional screw on other side). F) Screw to tighten anterior arch (along with additional screw on other side). G) Screw to lock height of bite bar (along with additional screw on other side). H) Screw to lock horizontal movement of bite bar (i.e., in/out of mouth). I) rod to secure body tube in place around mouse (along with rod on other side). J) Screw to secure adapter bar to headpost. K) Screw to secure adapter bar to the restraint (along with screw on other side).

The headpost-style restraint is also 3D printed using polycarbonate. It was inspired by a headpost designed by (Gutierrez-Barragan et al., 2022), but rather than permanently implanting the large ‘head-bar wing’, we implanted a small threaded collar on the skull to allow for the attachment of the ‘head-bar wing’ when the mouse is to be restrained. The motivation was to provide less of an impediment to the natural movement of the mouse in their cage when unrestrained. All designs have been made freely available ((https://doi.org/10.17605/OSF.IO/TYHCE).

### 2.4 Surgery

Mice in group 4 underwent surgery to have headposts implanted on their skulls. Mice were initially anesthetized with 4% isoflurane in oxygen and then switched to a 1.7 – 2 % mixture for maintenance. Once deeply anesthetized (confirmed using toe pinch), mice were injected with 0.05 mL/kg of Meloxicam to reduce postoperative pain and 0.05 mL/kg of Lidocaine to numb the surgical area. The hair on the scalp was removed with an electronic razor, eye ointment was applied, and a rectal thermometer was used to record temperature throughout the surgery. The mouse was placed in a stereotactic frame (71000 Automated Stereotaxic Instrument, RWD Life Science) and positioned with the skull level. After sterilizing the scalp, the skin above the skull was removed, and the soft tissue was cleared away. The skull was cleaned with saline and dried. The skull was then lightly scraped to provide a better surface for the bonding agent to adhere to. The bonding agent (All-Bond Universal, Bisco Dental) was applied to the skull and cured using an ultraviolet light. Dental cement (CoreFlo DC Lite, Bisco Dental) was subsequently applied, and the headpost was pressed onto the skull maintaining a high, constant pressure. The dental cement was cured, then the remaining exposed skull was covered with dental cement and cured. The mouse was then released from the stereotactic device and began a 10-day period of recovery prior to restraint acclimation. Additional injections of Meloxicam were given 24 and 48 hours after surgery.

### 2.5 Acclimation protocols

All mice were acclimated to the scanning environment in a mock scanner using either the non-invasive or headpost restraints (see 2.6). Three different acclimation protocols were used in the present study. Short (4-day; Group 1), medium (9-day; Groups 2 and 4) and long (13-day; Group 3) protocols were devised. The details for each protocol are shown in Table 2. These were designed to replicate protocols typically used in recent literature (refer to (Mandino et al., 2023)).

**Table 2.** Protocol used to acclimate each Group. Group 1 underwent the 4-day protocol (left), Group 2 and the headpost group underwent the 9-day protocol (middle) and Group 3 underwent the 13-day protocol (right).

| Day | Duration | Procedure |
| --- | --- | --- |
| 1 | 15 min | Manual handling (0 dB) |
| 2 | 30 min | Restrained (0 dB) |
| 3 | 45 min | Restrained (75 dB) |
| 4 | 1 hr | Restrained (100 dB) |
| 1 | 15 min | Manual handling (0 dB) |
| 2 | 10 min | Restrained (0 dB) |
| 3 | 20 min | Restrained (0 dB) |
| 4 | 30 min | Restrained (50 dB) |
| 5 | 45 min | Restrained (75 dB) |
| 6 | 1 hr | Restrained (100 dB) |
| 7 | 1 hr 15 min | Restrained (100 dB) |
| 8 | 1 hr 30 min | Restrained (100 dB) |
| 9 | 1 hr 30 min | Restrained (100 dB) |
| 1 | 15 min | Manual handling (0 dB) |
| 2 | 10 min | Restrained (0 dB) |
| 3 | 15 min | Restrained (0 dB) |
| 4 | 20 min | Restrained (50 dB) |
| 5 | 30 min | Restrained (75 dB) |
| 6 | 45 min | Restrained (75 dB) |
| 7 | 45 min | Restrained (75 dB) |
| 8 | 1 hr | Restrained (75 dB) |
| 9 | 1 hr | Restrained (75 dB) |
| 10 | 1 hr 15 min | Restrained (100 dB) |
| 11 | 1 hr 30 min | Restrained (100 dB) |
| 12 | 1 hr 30 min | Restrained (100 dB) |
| 13 | 1 hr 30 min | Restrained (100 dB) |

All acclimation sessions were started at approximately the same time each day to control for circadian rhythm phase. Day 1 consisted of the same procedure for Groups 1, 2, and 3. Specifically, mice were introduced to the restraint and allowed to freely explore it for the first few minutes. For the remaining duration of the session, mice would be encouraged to walk through the restraint. This involved the researcher picking up the mouse by the tail and guiding the subject into the apparatus. After successfully walking through the tube, a chocolate sprinkle would be provided as a reward. Subsequently, the subject would be repeatedly encouraged to walk through the restraint to establish that movement. Importantly, day 1 does not involve restraint. Each of the following sessions starts with this exercise for the first few minutes to encourage the intended behaviour.

Restraining the mouse using the proposed non-invasive restraint involves 3 steps: 1) broad restraint, 2) bite bar, and 3) fine restraint. Step 1 consists of letting the mouse walk through the body tube as performed previously but stopping the subject after their head passes through the neck plate and carefully lowering the neck plate into position. This effectively prevents the subject from moving further forward or backward in the restraint. To gradually expose mice to being fully restrained, Day 2 of each session consists of keeping the mouse in this coarsely restrained position only. Day 3 of acclimation onwards involves all 3 restraint steps. Step 2 of restraint uses the bite bar and anterior arch. This involves using the anterior arch to control the subject’s snout and guide the top front teeth (incisors) into the bite bar groove. Once in position, the bite bar and anterior arch are secured in place with fastening screws. Finally, step 3 is implemented consisting of the posterior arch being positioned over the posterior aspect of the subject’s head. Effectively, the neckplate and body tube prevent broad motion, and the combination of the bite bar, anterior arch and posterior arch prevent fine motion.

### 2.6 Mock scanner

It is common practice to design a “mock scanner” to simulate the scanning experience (Dinh et al., 2021; Lindhardt et al., 2022). This has the benefit of allowing for a more gradual acclimation as subjects can be exposed to restraint and MRI sounds independently without tying up the actual MRI scanner. Additionally, large numbers of animals can be acclimated at the same time. The mock scanner apparatus consists of a large box (65 cm wide, 52 cm long and 44 cm high) with PVC pipes of the same diameter as our gradient coil placed inside. A Bluetooth speaker is also placed in the box to play MRI sounds at designated sound levels (Table 2). These sounds were recorded directly from the MRI while performing the sequences used to scan subjects during acquisition. The audio was recorded and corresponding sound level was measured using a fibre optic microphone (OptiSLM 100, OptoAcoustics Ltd) placed inside the scanner at the same location as the subject. The maximum sound level of 91.6 dB was reached during the EPI fMRI scan. Acclimation session audio was played back at a maximum sound level of approximately 100 dB to ensure subjects were exposed to the maximum sound level they would experience in the scanner. Both the gradient-echo echo planar imaging (GE-EPI) sequence and anatomical Turbo RARE sequence were recorded. The audio clips were edited to create customized lengths such that each acclimation session included audio from both sequences of increasing lengths and sound levels (refer to Table *2*). Additionally, sound-suppression foam was adhered to the inner surfaces of the mock scanner box to reduce the noise heard from outside the acclimation room (i.e., to prevent other studies in nearby spaces from being affected). The foam is a multi-layer acoustic suppression medium consisting of a high-density rubber base (approximately 2 mm) with a foam backing (approximately 5 mm).

### 2.7 MRI acquisition

All acquisitions were performed at the Centre for Functional and Metabolic Mapping at The University of Western Ontario. MRI scans were acquired on a 9.4 T, 31-cm horizontal-bore magnet (Varian/Agilent, Yarnton, UK) with a 6-cm Magnex HD gradient insert (gradient strength=1000 mT/m, slew rate=9090 T/m/s) and Bruker BioSpec Avance NEO console running ParaVision 360 v3.3 (Bruker BioSpin Corp, Billerica, MA). An in-house 2-cm transmit/receive coil bent to conform to the mouse head curvature was used.

Groups 1, 2, and 3 had four 10-minute functional runs (two with a reversed phase encode direction) acquired per session (TR=1.5 s, TE=12 ms, resolution=0.3x0.3x0.4 mm^3^, slice gap=0.1 mm, matrix=64x32x30 voxels, 400 volumes, bandwidth=250 kHz). Following the functional scans, a T2-weighted Turbo RARE anatomical scan was acquired (TR=5.5 s, TE=30 ms, resolution=0.15x0.15x0.4 mm^3^, slice gap=0.1 mm, 192x96x30 voxels, flip angle=160 degrees). Group 4 was acquired previously and had slightly different acquisition parameters. Namely, 2 functional runs were acquired per session with 600 volumes each (15-min runs). All other parameters were the same.

### 2.8 fMRI analysis

All fMRI data were analyzed using the Rodent Automated Bold Improvement of EPI Sequences (RABIES version 0.5.1) (Desrosiers-Grégoire et al., 2024). All parameters used for RABIES processing stages are provided in Supplementary Table 1. Images were motion corrected using a trimmed mean across the acquired EPI frames in each run. They were then registered to a common space using an unbiased template as an intermediate step to improve registration (Avants et al., 2011). Image distortions due to susceptibility artifacts were corrected by nonlinearly registering EPI frames to the corresponding subject’s T2w anatomical image (Wang et al., 2017). This is performed after first applying a correction to all EPI frames to compensate for global static image intensity variation from the surface coil. Importantly, all transformations calculated during preprocessing (from motion correction to registration and susceptibility correction) are concatenated and applied in a single operation to avoid multiple stages of resampling. Final common space images were resampled to 0.2 mm x 0.2 mm x 0.2 mm prior to further processing.

Frame censoring was carried out using a framewise displacement (FD) threshold of 0.15 mm, i.e. half a physical voxel. This liberal threshold was chosen to deliberately include a degree of motion contamination in order to assess its impact on network quality. All frames exceeding this value, along with the preceding frame and the two following frames were removed (Power et al., 2012). If fewer than 200 volumes remained after censoring, the run was excluded from further analyses. Following this, data were detrended on a voxel-wise basis to remove linear drifts and converted to units of percent BOLD change through grand mean scaling (i.e., dividing timeseries by the grand mean signal – calculated over the whole brain – and multiplying by 100). The 6 motion parameters calculated during motion correction were used as regressors to further clean the data. Lastly, spatial smoothing (Abraham et al., 2014) was implemented using a full-width half maximum value of 0.3 mm.

### 2.9 Motion evaluation

Motion was estimated by registering EPI volumes and determining the displacements for each run. Although typical motion analyses involve measuring the 6 rigid body parameters, the voxel displacements between subsequent volumes are more important for data quality and better predict motion artifacts. This is because low frequency motion can be much more readily corrected for than high frequency spikes (Power et al., 2018; Soares et al., 2016). Here, we use the mean voxel-wise FD metric as performed in RABIES. It is calculated by taking the average of brain voxel displacements between each subsequent volume. A more detailed description of this metric can be found in the RABIES documentation (Desrosiers-Grégoire et al., 2024). This metric was used to compare motion levels between the 3 acclimatation protocols using the non-invasive restraint and the headpost restraint. The Kruskal-Wallis test was performed to determine significant differences followed by Bonferroni-corrected Wilcoxon rank sum test to determine which groups were significantly different.

Furthermore, to investigate which factors were most related to motion, correlation analysis was performed on subject age, number of times scanned (i.e. sessions), run number within a session, weight, and sex (Supplementary Figure 2). Each parameter value was determined at the time of each run, then Spearman’s rank correlation coefficient was determined between each parameter and the mean FD for the corresponding run.

### 2.10 Resting-state network analysis

Following preprocessing, fMRI data quality was analyzed to evaluate the performance of the two restraint approaches. Due to the subjective nature of evaluating the robustness of resting-state networks, we decided to compare our data using two previously derived components representing the well-established somatomotor network and default-mode network (DMN) provided with the RABIES software (Desrosiers-Grégoire et al., 2024). These consensus prior networks are based on 441 mice scanned with different awake and anesthesia protocols, at different field strengths and different sites. This was done by running dual-regression on our data using the prior component and then using Permutation Analysis of Linear Models (PALM) (Winkler et al., 2014) to obtain group statistical maps measuring the extent to which each group contained the prior component.

#### 2.10.1 Effect of acclimation protocol on network quality

First, we compared the network differences compared to the prior for our 4 awake groups. To ensure the same amount of data was used, the total number of frames acquired in the headpost group was calculated and the same quantity of data was randomly selected from each non-invasive group (*before* frame censoring). Group maps were then generated using PALM and z-scores were shown. To evaluate network quality, spatial specificity of the maps was measured using the Dice Similarity Coefficient (DSC) between the prior component with a z-threshold of 3.1 and the group map with a z-threshold of 3.0 for the somatomotor network and 3.1 for the DMN. The prior component threshold was chosen to represent a p value of 0.001 and the corresponding group map z threshold was selected to achieve the best overlap between the headpost group and the prior network. Multiple z values were tested to determine this optimal value for both networks. This same value was used for the non-invasive groups to fairly compare between groups. Additionally, the voxel-wise correspondence of z-values between the group and prior networks were obtained using the Pearson correlation coefficient (r) in the region of the network (by masking both prior and group networks by the voxels in the prior network that were greater or equal to a z-threshold of 3.1).

#### 2.10.2 Effect of motion and cumulative group acquisition time on network quality

Next, the effects of motion and cumulative group acquisition time were explored. Here, cumulative group acquisition time is defined as the total scan time after adding (concatenating) the scan time from all functional runs (for all subjects) used to make up the group map. The relative effects of motion and cumulative group acquisition time on network specificity and amplitude correspondence (i.e. the z-score of a voxel in the prior map vs the z-score of the same voxel in the restraint group maps) were both investigated. To do this, the non-invasive data from each acclimation protocol were first aggregated and then separated into 5 groups according to the amount of motion present, measured using the average run FD (namely motion level 1-5, Supplementary Table 2). Within each motion level, runs were randomly sampled to form groups with a total concatenated group acquisition time of 30, 60, 90, 120, 150, 180, and 210 minutes. This was accomplished through bootstrapping (randomly sampling the data without replacement). For each group, 10 random samples were selected, and PALM was used to attain group maps for both the somatomotor network and the DMN. Network specificity was again evaluated using the DSC of the group map and the prior component map. Similar to the acclimation group comparison, a z-value threshold of 3.1 was used for the prior component for the somatomotor network and DMN network (representing a p value of 0.001) and the z-value threshold of the somatomotor group network was 3.0 and the DMN group network was 3.1. Amplitude correlation was calculated by determining the correlation of voxel values between the prior network and corresponding group network (Figure *4* and Supplementary Figure 4). For each cumulative acquisition time, a one-way ANOVA followed by Tukey’s test for multiple comparisons was performed to determine the non-invasive motion groups that significantly differed from the headpost group.

The effects of cumulative group acquisition time and motion on network spatial specificity were further investigated by generalizing the relative relationships. All functional runs were aggregated and ranked by their average FD values, in ascending order of motion. Runs were grouped sequentially, starting with pairs, and the corresponding group map for each pair was generated. This map was then used to calculate the DSC. This process was repeated with progressively larger group sizes, including triplets, quartets, and so on, up to groups of ten. Linear trendlines were added to visualize the motion degradation effect for the different cumulative acquisition times (Supplementary Figure 3).

#### 2.10.3 Effect of motion censoring threshold on network quality

One of the most widely used and effective means of dealing with motion contamination is through implementing motion censoring (Power et al., 2012). Selecting too strict of a threshold will censor too much data and selecting too lenient of a threshold will allow too much degradation due to motion. To evaluate the impact of selecting different thresholds, somatomotor group maps for the different motion groups as outlined in Supplementary Table 2 were generated using censoring thresholds of 0.05, 0.10, 0.15 mm and without any threshold. DSC values between the group maps and prior map were determined and plotted in Supplementary Figure 5. A threshold of 0.15 mm, corresponding to half a physical voxel (in-plane) was selected as optimum.

#### 2.10.4 Effect of data density per subject on network quality

Study design requires the selection of the number of subjects to use and how much data to acquire per subject. To evaluate the effect of this choice on network quality, 20 runs from one subject (resulting in a cumulative acquisition time of 200 min) was used to produce a group somatomotor map and compared with a group comprised of a single run from 20 different subjects (also 200 min). DSC and amplitude correlation values were calculated and compared between them. Results are illustrated in Supplementary Figure 7.

### 2.11 Stress response

Animal weight was measured for all mice throughout acclimation and scanning. Real-time video monitoring during scans was done using an MR-compatible camera acquiring frames at 20 Hz (“12M-I *newSensor*” with integrated LED light, MRC Systems GmbH, Germany). A subset of mice (n = 21) was used to quantitatively assess CORT changes throughout the short (4-day), medium (9-day) and long (13-day) acclimation sessions. In contrast to the conventional extraction of CORT through blood serum which may cause a confounding stress response (Eleftheriou et al., 2020; Rowland & Toth, 2019; Russo et al., 2021), we employed extraction from fecal pellets similar to (Russo et al., 2021). This was performed by collecting the fecal pellets defecated during each session. CORT was extracted and the concentrations were determined using an ELISA following the supplier’s manual (Arbor Assays DetectX Corticosterone Chemiluminescent Immunoassay Kit, catalog number K014-C). All concentrations were measured in duplicate and are expressed as ng of CORT per mg of fecal matter.

CORT was analyzed on days 1, 2, and 4 for the 4-day protocol, days 1, 2, 4, and 9 for the 9-day protocol, and days 1, 2, 4, 9, and 13 for the 13-day protocol. All samples were read from a luminescence-capable plate reader (SpectraMax® M5 Multimode Plate Reader, Molecular Devices, USA). One-way ANOVA was used to compare CORT concentrations in each acclimation group. No CORT analysis was performed on the headpost group as it was a dataset acquired previously for a separate study. However, CORT data from head fixed animals using similar headposts has been obtained by others (Lindhardt et al., 2022).

## 3 Results

### 3.1 Motion evaluation

We first analysed scan motion levels to determine if motion depends on the acclimation protocol used (Figure *2*). The FD for each voxel was measured and the average across each run was computed. Figure *2* (a) shows a representative motion plot with the 6 rigid body parameters. The corresponding mean voxel-wise displacements are shown below in Figure *2* (a). A histogram is plotted in Figure *2* (b) showing all FD measures for all acquired volumes. This metric was then used in Figure *2* (c) to compare motion between cohorts. Motion was significantly lower in the headpost group than all other groups. Note, after censoring, 18.9% of the total number of acquired volumes were censored with 18 of the 236 runs (7.6%) excluded due to being left with fewer than 200 volumes after censoring. A comprehensive comparison of the censored volumes based on group can be found in Supplementary Table 2.

**Figure 2.**
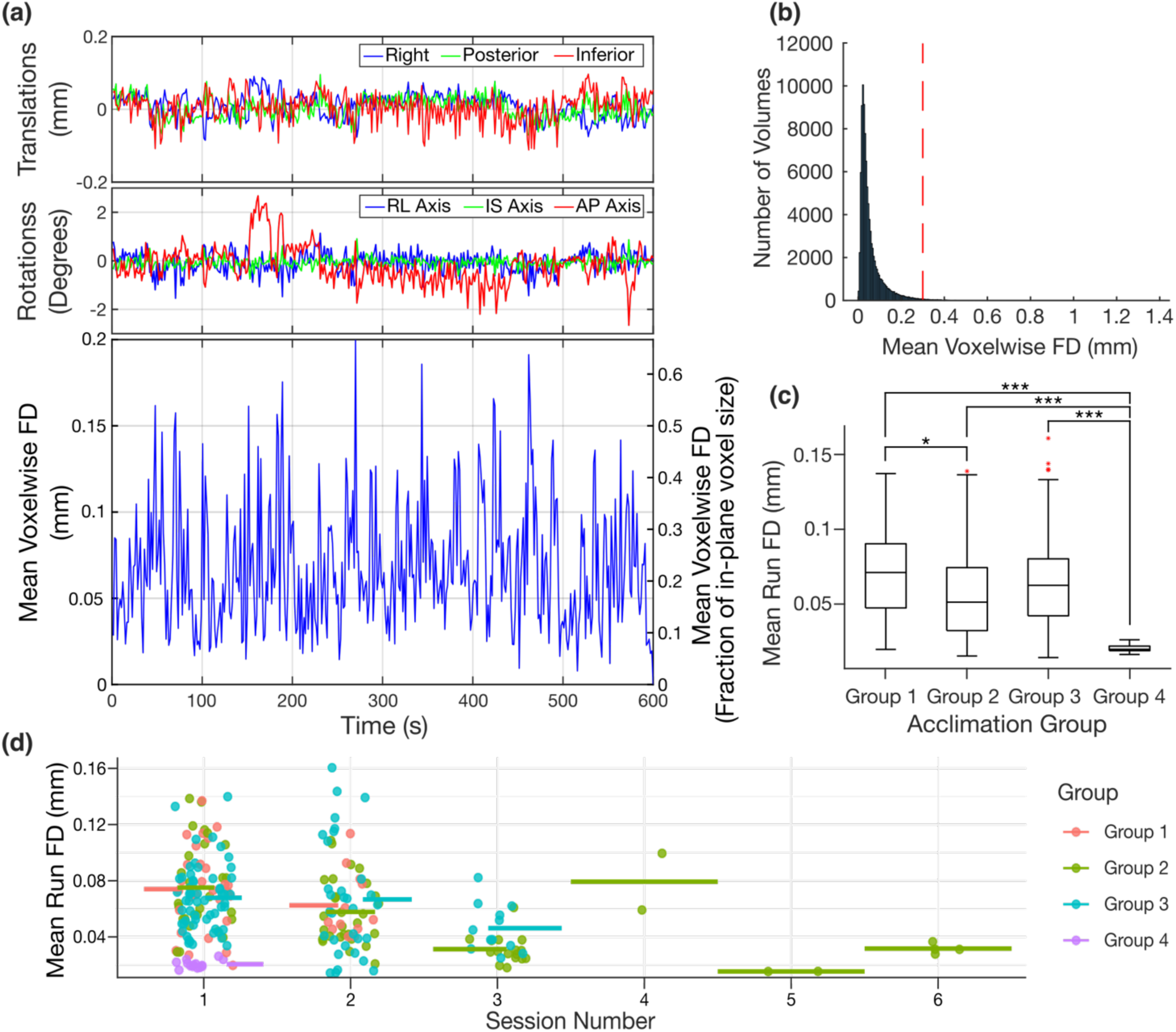
Scan motion levels for the different restraints and acclimation protocols. (a) Representative plot displaying the motion parameters (3 translations – top, 3 rotations – middle, and the corresponding mean voxel-wise framewise displacements (FD) – bottom). The plot is expressed in FD magnitude (left y-axis) and as a fraction of the in-plane voxel size (0.3 mm – right y-axis). (b) A histogram displaying mean voxel-wise FD values for all volumes for all subjects. The red vertical line indicates the in-plane voxel size (0.3 mm) for reference. (c) Boxplots comparing the motion between the four cohorts prior to censoring. Outliers are shown as red ‘*’. The mean run voxel-wise FD values are calculated by taking the average of all brain voxel displacements between subsequent volumes (* = p < 0.05, ** = p < 0.01, *** = p < 0.001, Kruskal-Wallis rank sum test followed by Bonferroni-corrected Wilcoxon rank sum test). (d) Mean run FD values as a function of session number. Colours differentiate the acclimation groups. The horizontal bars represent the mean FD for each session and are coloured according to their corresponding group. Group 1 = non-invasive mice acclimated using 4-day protocol, Group 2 = non-invasive mice acclimated using the 9-day protocol, Group 3 = non-invasive mice acclimated using the 13-day protocol, Group 4 = headpost mice acclimated using the 9-day protocol.

### 3.2 Resting-state networks

To avoid the bias of using components generated from this dataset and promote standardization across centres, dual regression was performed using previously generated components through independent component analysis that established well-known resting-state networks (Desrosiers-Grégoire et al., 2024). To confirm the networks were recoverable in our datasets, dual regression of the somatomotor network and DMN were performed for each scan and data were grouped by acclimation group (Figure *3*). The DSC representing the amount of overlap and correlation coefficient representing the amplitude correspondence between the networks is overlaid above each network. Note, only a sample slice is shown however the DSC and correlation coefficients were calculated from the entire volume.

**Figure 3.**
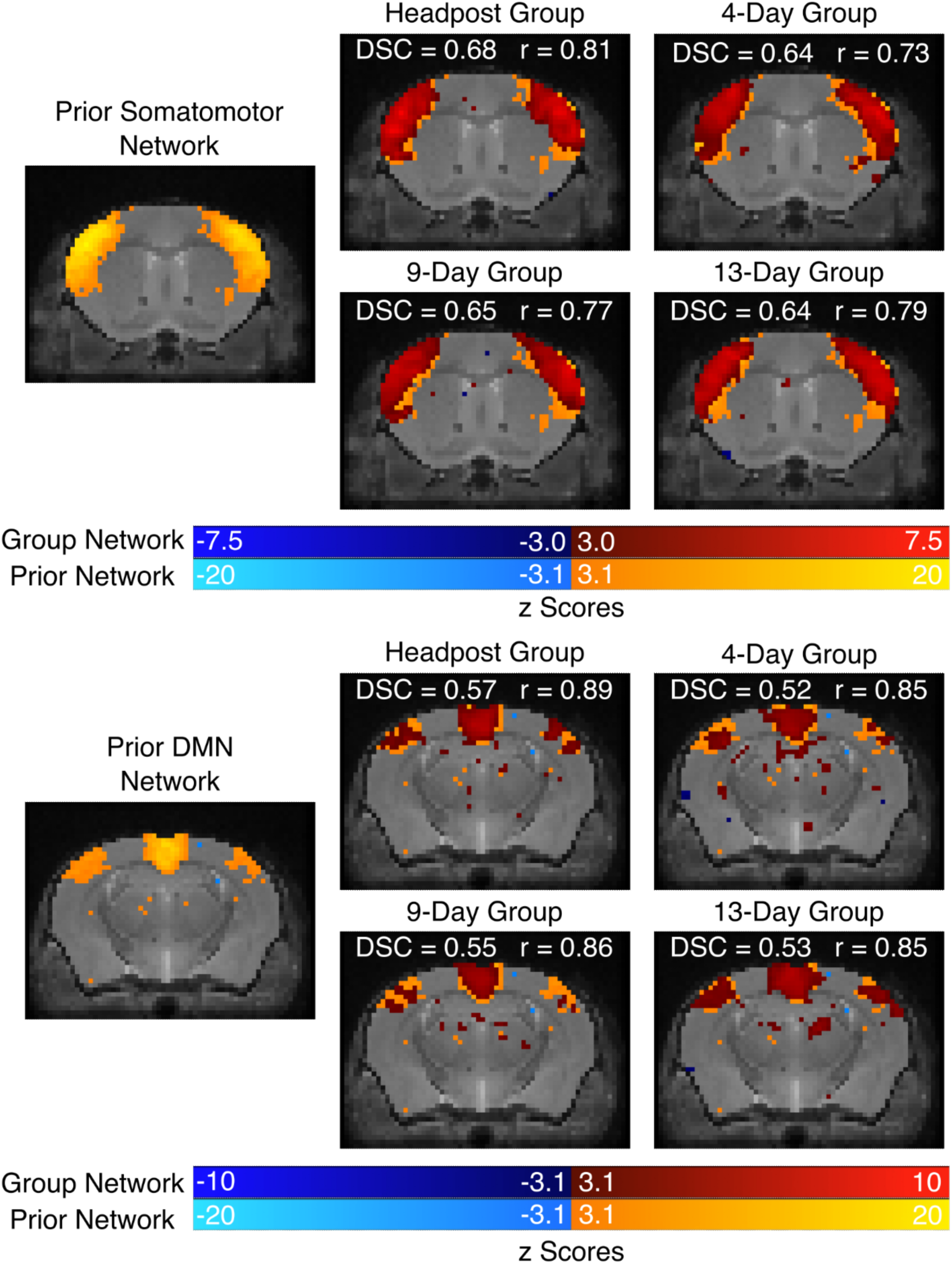
Network quality comparison of the 3 non-invasive protocols and the headpost group using the Dice Similarity Coefficient (DSC) and amplitude correspondence (using the Pearson correlation coefficient, r) with the externally derived somatomotor (top) and DMN (bottom) components *(Desrosiers-Grégoire et al., 2024)*. Note, DSC and correlation values were calculated using the 3D volumes (prior components and groups components) but only one sample slice is shown for simplicity. The network maps shown on the left are the externally derived components. The remaining images show the group networks (dark blue – dark red colour bar) overlayed over the prior network (light blue – light orange colour bar). Corresponding DSC and r values are annotated over each network.

We further investigated the relationship between motion, sample size and network quality by looking at how varying levels of motion and cumulative group acquisition time affect quality (i.e., DSC and amplitude correlation) relative to the prior network component (Figure *4*). Supplementary Figure 4 shows the resulting correlation plots for a single sample from each motion level and cumulative group acquisition time. Past 120 min of acquired data, both DSC and amplitude correlation analyses show clear statistical differences between headpost and non-invasive groups. Supplementary Figure 3 shows the effect of motion on spatial specificity for different cumulative group acquisition times.

**Figure 4.**
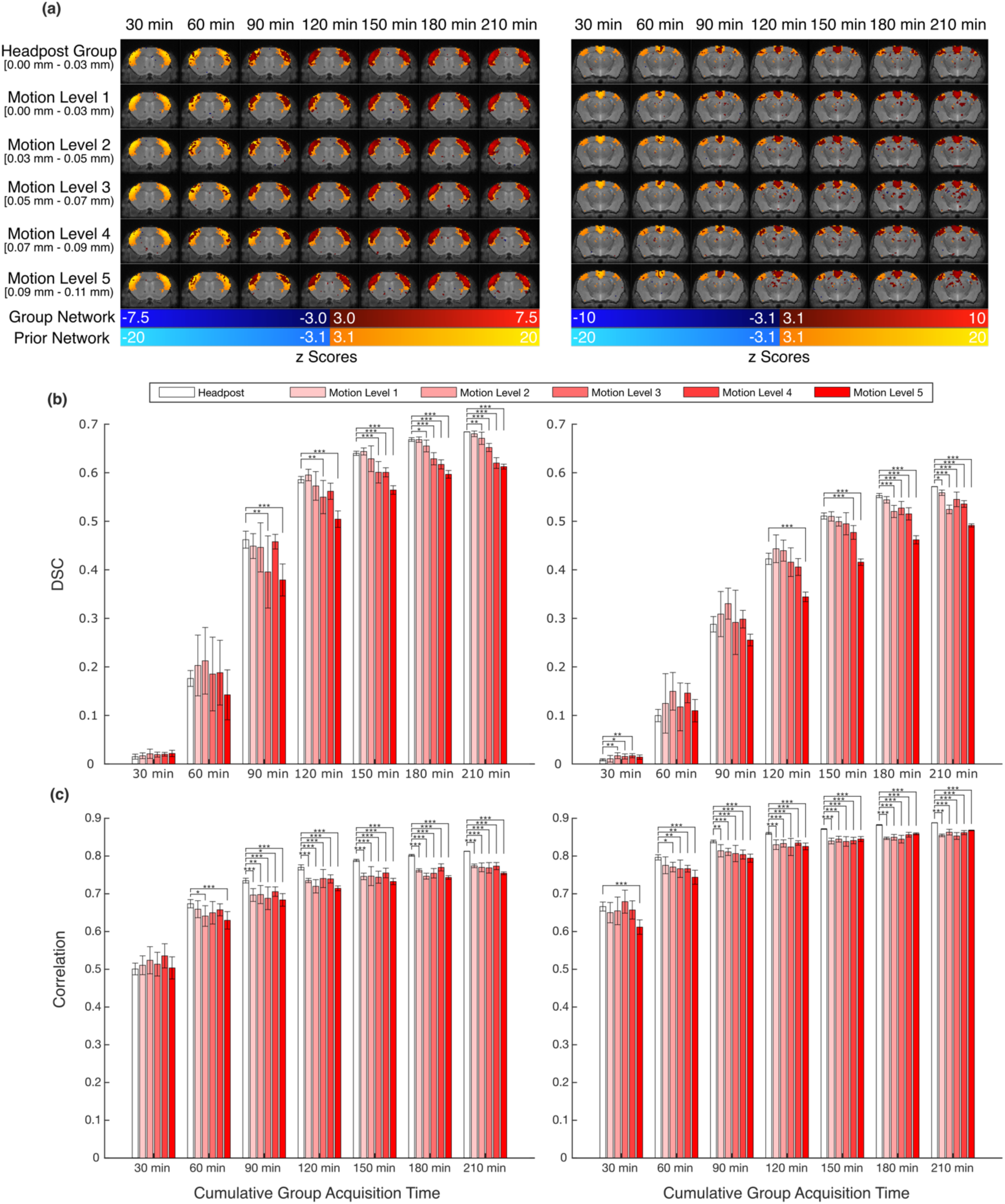
Group network quality assessment for the headpost and non-invasive data relative to the prior network component. Motion groups (i.e., Motion Levels 1-5) were determined by concatenating all non-invasive data and regrouping runs based on their average framewise displacement (FD). Motion Level 1: 0-0.03 mm, Motion Level 2: 0.03 – 0.05 mm, Motion Level 3: 0.05 – 0.07, Motion Level 4: 0.07 – 0.09, and Motion Level 5: 0.09 – 0.11. Varying numbers of runs were selected from each group to create samples with cumulative pre-censoring acquisition times of 30, 60, 90, 120, 150, 180 or 210 minutes. (a) Sample slices of the z-maps of the somatomotor prior component (left) and DMN prior component (right) illustrated in light orange/light blue and the corresponding group network overlayed in dark red/dark blue. The range of average FD values for each group are displayed below each Motion Level on the left (square brackets indicate inclusivity, parenthesis indicate non-inclusivity). (b) Corresponding Dice Similarity Coefficient (DSC) values to the network overlaps shown in (a) for the somatomotor network (left) and DMN (right). (c) Correspondence of z-values compared voxel-wise between the prior map and the group map masked to include voxels in the prior network region for the somatomotor network (left) and DMN (right). Error bars in (b) and (c) represent the standard deviation of 10 random samples selected from the respective group. Groups that significantly differed from the headpost group for each cumulative acquisition time are shown (* = p < 0.05, ** = p < 0.01, *** = p < 0.001, for Tukey’s test for multiple comparisons following one-way ANOVA).

### 3.3 Stress

Mouse stress was approximated using weight loss and fecal CORT concentration over the course of the acclimation sessions. Weight was measured each day of acclimation. Values were normalized to the weight on day 1 and plotted for each cohort separately (Figure *5* (a) and (b), Supplementary Figure 6 (a) and (b)). Paired t-tests were conducted to test for statistical differences between first and last days of acclimation for each group. Headpost (males), 9-day and 13-day groups all had significant weight losses. Fecal corticosterone metabolite (FCM) was measured from a subset of mice from each non-invasive group. Results are shown in Figure *5* (c) and Supplementary Figure 6 (c) and (d). Note, the headpost group was acquired previously and fecal samples were not collected. One-way ANOVA was conducted. Paired t-tests could not be performed since several groups had missing samples (due to the subject not defecating during the respective acclimation session). There was no significant difference in CORT levels between sampling days across the acclimation protocols.

**Figure 5.**
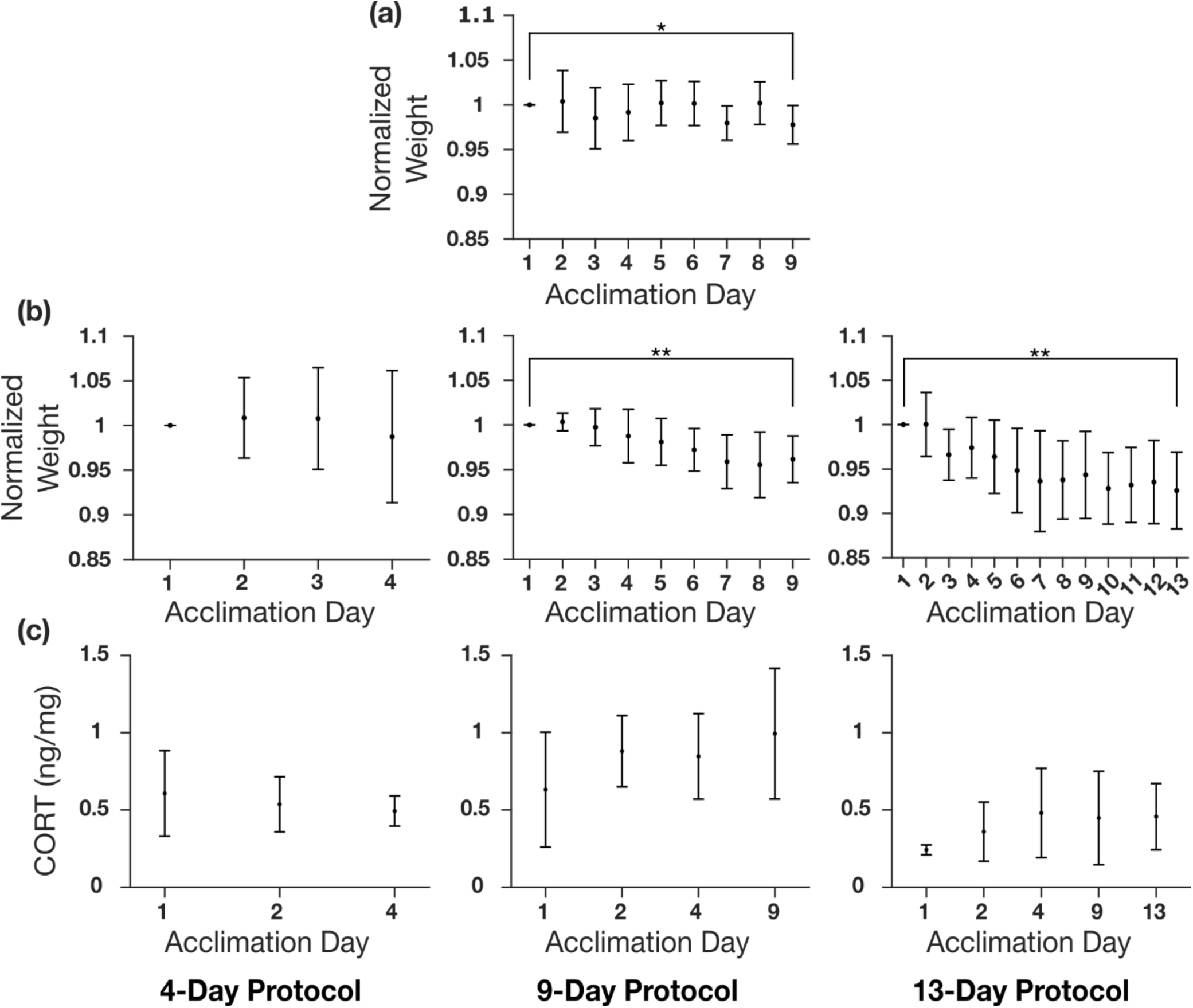
Weight and fecal corticosterone (CORT) changes across acclimation sessions. (a) Weight changes for all subjects (n = 7) in the headpost group normalized to the weight on day 1. The weight on the last day was significantly less than the first day (paired t-test, p < 0.05). (b) Weight changes of all subjects in the 4-day, 9-day and 13-day groups over the course of the different acclimation lengths normalized to the weight on day 1. Note the headpost group underwent the same 9-day acclimation session as the 9-day non-invasive group. Both the 9-day group and 13-day group showed significant weight decreases (paired t-test, p < 0.01). (c) CORT concentration changes over the course of the three acclimation durations. Note, the headpost group was acquired in a separate study and CORT samples were not collected. There were no significant CORT changes across acclimation sessions (one-way ANOVA, p > 0.05).

## 4 Discussion

All subjects used in this study were successfully acclimated to either the headpost or non-invasive restraint using the described protocols. The new non-invasive restraint contains several adjustable features to tailor the fit for each mouse regardless of size (i.e., 16 – 37 g as tested here). Similar to previous non-invasive restraints (Madularu et al., 2017), our restraint is open source, 3D printed and modular in nature, allowing for easy modification by other labs. Furthermore, no anesthesia is required for setup and experienced handlers can position acclimated mice in the restraint device and insert into the scanner in less than 5 minutes.

Three different acclimation procedures were explored representing a range of protocols typically found in recent awake mouse MRI literature (see (Mandino et al., 2023; Zeng et al., 2022) for comprehensive acclimation protocol comparisons). Based on the motion analysis, headpost mice moved consistently the least, as seen in Figure *2* (c). Individual run average FD measures and FD peaks can be seen in Supplementary Figure 1 (a) and (b), respectively. The 4-day group had the largest fraction of volumes censored and largest fraction of runs excluded due to being above the motion threshold (Supplementary Table 2). Even so, the network specificity between the 4-day, 9-day and 13-day groups were almost identical (Figure *3*). For both networks, all non-invasive groups were less than 10% lower in terms of spatial specificity and amplitude correlation compared to the headpost group. Clearly the headpost outperforms the non-invasive restraint, but the data also show that this can be compensated by using either slightly larger groups and/or marginally longer acquisition times.

Some groups have implemented more realistic acclimation procedures by performing sessions in the MRI scanner itself rather than a mock scanner (Dinh et al., 2021). This exposes mice to the exact environment they will experience during scans which helps with habituation. This has downsides in terms of throughput (1 animal at a time), the cost of scanner time and limiting scanner access for other studies in a multi-user facility. To evaluate the potential additional benefit of such an approach, we analyzed the movement (in terms of average FD) as a function of the scan session number (Figure 2 (d), Supplementary Figure 1 (a)). This analysis showed that mice do move marginally less with additional scanning sessions. We also found that motion was less in older animals. Motion as a function of weight, sex and run number within a session was also evaluated, though no correlation was found (Supplementary Figure 2). Our results suggest that minor improvements in motion control are to be expected as the animal acclimates to the actual environment of the MRI scanner despite the pretraining protocols.

The censor threshold we selected was 0.15 mm. This liberal threshold (half a voxel) allows for higher-motion volumes to remain and was important to evaluate the effect of these motion-contaminated volumes in network quality. Figure *4* outlines these results for the robust somatomotor network and the less-prominent DMN. To identify the impact of selecting stricter censor thresholds, a single sample from each group in Figure 4 from the somatomotor network analysis was evaluated with censor thresholds of 0.05 mm, 0.10 mm, and without a threshold (Supplementary Table 3 and Supplementary Figure 5). The choice of threshold has very little effect on spatial specificity for low-motion groups. With high-motion groups, stricter thresholds are correlated with lower spatial specificities due to higher data attrition. This justifies the selection of the 0.15 mm censor threshold and supports the claim that the cumulative group acquisition time better predicts group network quality compared to the level of motion contamination (at least for the levels of motion contamination seen here).

Non-invasive restraints are more vulnerable to residual motion compared to headpost restraints, since the latter directly fixates the skull. However, we have found the impact of the motion confound reduces tremendously with increased sample size as demonstrated in Figure *4*. Since motion tends to affect a relatively constant proportion of data, an increased sample size will allow for increased statistical power after censoring of motion-affected frames. Even at the extreme end, in untrained young human subjects, Smith and colleagues evaluated how harmful this motion is to data quality using similar metrics outlined here. In the end, they conclude a conservative design would be to acquire 2 to 3 times more data than would be expected for the same study without high-motion subjects (Smith et al., 2022). The present study suggests a much lower quantity of data is required. Despite allowing more than 3x the motion than the headpost cohort (Figure *2* (c)), acquiring 25% more data with the non-invasive restraint will compensate for censored volumes assuming most motion is at the high end allowed by the restraint. As stated previously, we have also found an inverse relationship between FD and the number of sessions the mice undergo (r = -0.42, Supplementary Figure 2). In combination with Figure 2 (d), this demonstrates that as a mouse gets scanned more often, there is a subtle trend to move less. Figure 2 (d) also shows that there is no substantial difference between Groups 1-3 during the first two scan sessions for which we have data from all groups. This suggests that a 4 day acclimation period is sufficient before starting the MRI scans. Though, researchers may find it beneficial to increase the number of scanning sessions per subject as this will result in less motion during later scans due to further acclimation to the scanning environment and consequently better data quality.

The effects of motion are only seen past 50 min of cumulative group acquisition time. From Supplementary Figure 3, this is evident from the emergence of the negative slope of the linear trendlines. The slope only becomes noticeable in the 50 min group and above. The groups with less cumulative acquisition time contain other sources of variation such that the effects of motion are not apparent. Individual network variability can cause such variation (Bergmann et al., 2020; Fu et al., 2023). In Figure *4* the effects of motion are only seen at 90 min for the somatomotor networks and 120 min for the DMN as can be seen by the significantly lower spatial specificity for higher motion levels. Note, however that the difference between the low motion headpost group and the low motion non-invasive groups (i.e., Motion Level 1 and Motion Level 2) only becomes apparent at higher cumulative group acquisition times (i.e., >150 min) when the other sources of variation are presumably averaged out.

Further comparing the robust somatomotor network with the less prominent DMN, a few more false activations can be seen in the DMN. This is quantified by the lower DSC values compared to the somatomotor network. The same relationship is not seen with amplitude correlation. In fact, the DMN has a higher amplitude correlation indicating a more similar spatial distribution of voxel magnitudes with the prior component. This can be explained by the increased homogeneity of the DMN compared to the somatomotor network. Still, the relationship between network quality and increased group acquisition time is apparent for both networks (Figure *4*) as seen by the rising network quality with increasing cumulative group acquisition times for all motion level groups. To investigate whether these false activations are in fact true activations seen in awake datasets without the anesthesia confound (compared to the prior generated by predominantly anesthetized data), an open-source dataset was downloaded (Yun & Grandjean, 2020) and processed using the same pipeline in this manuscript. This dataset contains mice scanned awake, anesthetized using isoflurane, and anesthetized using both medetomidine and isoflurane. The resulting DMN group maps were compared and illustrated in Supplementary Figure 8. All groups show similar activation in subcortical regions not active in the prior network, indicating these regions are likely due to noise rather than being dependent on anesthesia use.

Sample size is a current topic of controversy among animal imaging researchers (Grandjean et al., 2024; Mandino et al., 2023). There is motivation from an ethics and cost perspective to use smaller sample sizes, whereas from a statistical perspective, large sample sizes are preferred. Fortunately, here the slightly higher motion corrupted data benefits the same amount from increased group cumulative acquisition time compared to the headpost restraint (Figure *4*, Supplementary Figure 3). For instance, all groups have a spatial specificity above 0.4 (Figure *4* (b)) with 120 min of cumulative acquisition time for the somatomotor network and with 150 min for the DMN. Similarly, this amount of data gives an amplitude correspondence greater than 0.7 for all motion groups for the somatomotor network, and above 0.8 for the DMN (Figure *4* (c)). These results indicate the amount of data is much more important than the amount of motion during the acquisition. Indeed, with long cumulative acquisition times (approaching 210 min), the spatial specificity difference between motion levels seems to be solely based on the number of frames censored. It is possible higher amounts of motion (i.e., greater than the in-plane voxel size) would further degrade the networks, but that was not investigated here as almost all (98.8%) displacements were under the in-plane voxel size (Figure *2* (b)). Surprisingly, we do not see this same relationship when looking at the amplitude correspondence (Figure 4 (c)). Approaching higher cumulative group acquisition times, the headpost group emerges with a statistically significant but marginal increase over all non-invasive motion level groups, however no clear relationship emerges among motion level groups like we see in the spatial specificity analyses. This suggests that this amplitude correspondence metric is independent of motion, at least at the levels observed here. However, increased cumulative acquisition time still increases correlation values of all groups indicating differences between the headpost and non-invasive restraint can be addressed by simply acquiring more data. This increase in data can come either from adding more sessions with the same subjects or from keeping the number of sessions constant while including additional subjects. In both cases, the effect on network quality is comparable (Supplementary Figure 7). However, scanning the same participants multiple times may be more advantageous, as it also enhances acclimation for later sessions, as stated previously.

Stress has been shown to be a critical measure for awake rodent fMRI as it has been shown to impact functional connectivity (Grandjean et al., 2016; Maron-Katz et al., 2016). Indeed, the MRI environment presents several stress-inducing factors including physical restraint in a dark, loud, and enclosed environment, with very loud noises. The acclimation procedure outlined here effectively habituates mice to these factors except for the magnetic field. Nonetheless, all animals in all acclimation groups show a modest decrease in mean FD with each subsequent session in the MRI, suggesting that there are environmental differences between the mock scanner and the real scanner.

There is a statistically significant weight loss between the first and last days of acclimation in the headpost group, and the 9-day and 13-day non-invasive groups (Figure *5* and Supplementary Figure 6). Although no loss is observed in the 4-day group, this is likely because the 4-day duration is too short to observe the change. While these weight losses are statistically significant (paired t-test, p < 0.05), they remain below 15%, which is the threshold considered a concern for animal welfare by most animal care committees.

FCM was measured and compared for several selected days throughout each non-invasive acclimation group (Figure *5* and Supplementary Figure 6). Specifically, fecal samples were used over serum samples to minimize unrelated stress to the animals (Kroll et al., 2021). For the same reason, day 1 was selected as our baseline as it is well known that CORT is only detectable in feces hours after the stressful event (Rowland & Toth, 2019). It is also sensible to assume the FCM measurement on Day 1 is in fact a baseline measurement because 1) any CORT would not be detectable within 20 min (maximum duration between stress event onset and fecal excretion) of the stressor, and 2) the procedure on day 1 is the least stressful and likely would not result in any significant increase in CORT (e.g., Lindhardt et al. demonstrated that even a more intensive protocol involving 20 minutes of restraint on Day 1 did not result in a significant increase in CORT (Lindhardt et al., 2022)).

Surprisingly, no significant change in CORT concentration was observed in any of the acclimation groups. Consequently, these results would indicate no relationship between acclimation session and stress in mice and stress would not explain the observed difference in motion between the 4-day and 9-day acclimation groups. This is an unexpected result compared to the literature (Ferris et al., 2011; Gutierrez-Barragan et al., 2022; Lindhardt et al., 2022; Low et al., 2016). This could be due to the measurement of FCM rather than more acutely sensitive measures of CORT such as through plasma samples. FCM is known to integrate the CORT levels over relatively long periods of time (hours to days) (Rowland & Toth, 2019). It is possible our maximum 1.5-hour stressor did not produce a detectable CORT increase over the 48-hour integration time. (Lindhardt et al., 2022) used FCM and did find a difference on day 11 of acclimation compared to day 1. However, they used a more aggressive acclimation protocol than used here, therefore it is possible the animals showed a larger stress increase on the first few days. Those animals also underwent surgery which imposes an additional variable to consider.

Due to the results of this study, along with indications from related literature, the authors recommend future CORT experiments be acquired over longer durations each session. That is to say that single time-point measurements of CORT can be misleading and are difficult to interpret. Further, it should be noted that it is still not fully understood what increases in CORT represent and indeed, CORT levels naturally vary widely dependent on many factors (Rowland & Toth, 2019). While it is generally thought that increases correspond with increases in negative stress, it has also been postulated that elevations in hypothalamic–pituitary–adrenal (HPA) axis and basal glucocorticoid levels may be linked with stress adaptation, a positive outcome rather than excess stress (Rowland & Toth, 2019). More stress-targeted studies are required to both characterize and better understand the relationship of CORT concentration on perceived stress.

### 4.1 Limitations

Like all T2*-weighted sequences, care should be taken to minimize the effect of all major sources of susceptibility artifacts. This includes avoiding the use of magnetic materials and avoid pockets of air near tissue. One such source relevant to the proposed restraint was the susceptibility artifact caused by the ears of the mouse. Since the posterior arch can fold the ears onto the head, care should be taken to ensure the ears are folded towards the posterior and lateral regions of the head rather than anterior and medial.

Although the restraint is designed to allow for some tasks to be readily performed, task-based imaging was not conducted in the current study. It can be postulated that tasks may cause task-induced motion, the effects of which are amplified in restraints allowing for increased baseline levels of motion. However, since resting-state networks were clearly identifiable, it is likely task-based networks will only be clearer due to the higher activation and statistical power associated with task-based protocols (Zhao et al., 2023).

An additional limitation of this work is it does not consider intra-volume motion. Consequently, acquisitions with large movements that occur and return to the original position during the same TR may not be robustly corrected and can also have degrading effects (Kim et al., 1999) which may explain some of the variance observed in the network quality analyses.

Lastly, this study focused on the network measures using dual-regression of group ICA-based analyses. Although this is a very common approach used in resting-state fMRI studies, investigation into the effects of motion and cumulative group data quantity on other resting-state analysis approaches including seed-based and connectivity matrix-based approaches were not conducted.

## 5 Conclusions

An open-source, non-invasive restraint was developed and evaluated along with three different acclimation protocols. After evaluating motion and network quality using two well-established brain networks generated from a variety of large-N multi-site studies, all acclimation protocols were found to have similar spatial similarity and amplitude correlation values, with very small improvements as the acclimation period increased from 4 days to 13 days. Very small improvements in FD were observed with each session in the MRI independent of acclimation period. Our results suggest that 4 days of acclimation are sufficient to introduce an animal to the MRI, and that useful data can be acquired even in the first MRI session. Thus, the first MRI session serves in part as a further acclimation session, but still allows for the acquisition of high-quality data. Although the mean FD for the non-invasive restraint was 2-4x that of the headpost animals, we observed little difference between both restraints using our metrics of spatial specificity and amplitude correlation. Our analysis showed that by acquiring approximately 25% more data to conservatively compensate for censoring of high motion frames, the non-invasive restraint can be employed to achieve group resting state networks at comparable quality to headpost approaches.

Surgical headpost restraints have pushed the mouse fMRI field forward by allowing for awake scanning with very little motion. However, certain situations warrant the need for awake approaches that require no surgery or administration of even single doses of anesthesia. If these considerations are important for a study, our non-invasive awake MRI method provides an elegant solution. The method has the potential to broaden the scope of possible awake mouse fMRI studies, with future applications providing insights into cognition and the fundamental basis of brain function.

## Data and Code Availability

All data and code have been made freely available (10.17605/OSF.IO/TYHCE).

## Author Contributions

S.L., and R.M. designed the study, S.L., A.E., M.B., and M.N. collected and analyzed the data, S.L., K.M.G., P.Z., and R.M. designed the non-invasive restraint, S.L. wrote the initial draft of the manuscript with feedback and review from all authors.

Funding

This work was supported by funding from the Canadian Institutes for Health Research (FDN-148453), a Brain Canada Platform Support Grant and a Canada First Research Excellence Fund award to BrainsCAN and a New Frontiers in Research Fund Transformation award (NFRFT-2022-00051).

## Declaration of Competing Interests

Nothing to declare.

## Acknowledgements

We thank Alex Li for his help setting up and troubleshooting MRI scans; Gabriel Desrosiers-Grégoire, Gabriel A. Devenyi, and Mallar Chakravarty from The Douglas Research Centre for technical assistance with RABIES; and the Animal Care Committee from The University of Western Ontario and veterinarians for feedback and suggestions while developing the non-invasive restraint.

## Notes

https://doi.org/10.17605/OSF.IO/TYHCE

### Competing Interest Statement

The authors have declared no competing interest.

### Summary of Updates

A few additions to the Methods and Discussion; Two additional supplementary figures.

## References

Abraham, A., Pedregosa, F., Eickenberg, M., Gervais, P., Mueller, A., Kossaifi, J., Gramfort, A., Thirion, B. & Varoquaux, G. (2014). Machine learning for neuroimaging with scikit-learn. Frontiers in Neuroinformatics, 8, 14. 10.3389/fninf.2014.00014

Avants, B. B., Tustison, N. J., Song, G., Cook, P. A., Klein, A. & Gee, J. C. (2011). A reproducible evaluation of ANTs similarity metric performance in brain image registration. NeuroImage, 54(3), 2033–2044. 10.1016/j.neuroimage.2010.09.025

Bergmann, E., Gofman, X., Kavushansky, A. & Kahn, I. (2020). Individual variability in functional connectivity architecture of the mouse brain. Communications Biology, 3(1), 738. 10.1038/s42003-020-01472-5

Bianchi, S. L., Tran, T., Liu, C., Lin, S., Li, Y., Keller, J. M., Eckenhoff, R. G. & Eckenhoff, M. F. (2008). Brain and behavior changes in 12-month-old Tg2576 and nontransgenic mice exposed to anesthetics. Neurobiology of Aging, 29(7), 1002–1010. 10.1016/j.neurobiolaging.2007.02.009

Chen, X., Tong, C., Han, Z., Zhang, K., Bo, B., Feng, Y. & Liang, Z. (2020). Sensory evoked fMRI paradigms in awake mice. NeuroImage, 204, 116242. 10.1016/j.neuroimage.2019.116242

Desai, M., Kahn, I., Knoblich, U., Bernstein, J., Atallah, H., Yang, A., Kopell, N., Buckner, R. L., Graybiel, A. M., Moore, C. I. & Boyden, E. S. (2011). Mapping brain networks in awake mice using combined optical neural control and fMRI. Journal of Neurophysiology, 105(3), 1393–1405. 10.1152/jn.00828.2010

Desjardins, M., Kılıç, K., Thunemann, M., Mateo, C., Holland, D., Ferri, C. G. L., Cremonesi, J. A., Li, B., Cheng, Q., Weldy, K. L., Saisan, P. A., Kleinfeld, D., Komiyama, T., Liu, T. T., Bussell, R., Wong, E. C., Scadeng, M., Dunn, A. K., Boas, D. A., … Devor, A. (2019). Awake Mouse Imaging: From Two-Photon Microscopy to Blood Oxygen Level–Dependent Functional Magnetic Resonance Imaging. Biological Psychiatry: Cognitive Neuroscience and Neuroimaging, 4(6), 533–542. 10.1016/j.bpsc.2018.12.002

Desrosiers-Grégoire, G., Devenyi, G. A., Grandjean, J. & Chakravarty, M. M. (2024). A standardized image processing and data quality platform for rodent fMRI. Nature Communications, 15(1), 6708. 10.1038/s41467-024-50826-8

Dinh, T. N. A., Jung, W. B., Shim, H.-J. & Kim, S.-G. (2021). Characteristics of fMRI responses to visual stimulation in anesthetized vs. awake mice. NeuroImage, 226, 117542. 10.1016/j.neuroimage.2020.117542

Dumont, J. R., Sheppard, P. A. S., Fodor, C., Coto, M. A., Yang, S., Saito, T., Saido, T. C., Rylett, R. J., Prado, M. A. M., Bussey, T. J., Saksida, L. M. & Prado, V. F. (2025). Impaired Cognitive Flexibility With Preserved Learning in an Amyloid Precursor Protein Knock-In Mouse Model of Amyloidopathy. *Genes*, Brain and Behavior, 24(3), e70024. 10.1111/gbb.70024

Eleftheriou, A., Palme, R. & Boonstra, R. (2020). Assessment of the Stress Response in North American Deermice: Laboratory and Field Validation of Two Enzyme Immunoassays for Fecal Corticosterone Metabolites. Animals, 10(7), 1120. 10.3390/ani10071120

Fadel, L. C., Patel, I. V., Romero, J., Tan, I.-C., Kesler, S. R., Rao, V., Subasinghe, S. A. A. S., Ray, R. S., Yustein, J. T., Allen, M. J., Gibson, B. W., Verlinden, J. J., Fayn, S., Ruggiero, N., Ortiz, C., Hipskind, E., Feng, A., Iheanacho, C., Wang, A. & Pautler, R. G. (2022). A Mouse Holder for Awake Functional Imaging in Unanesthetized Mice: Applications in 31P Spectroscopy, Manganese-Enhanced Magnetic Resonance Imaging Studies, and Resting-State Functional Magnetic Resonance Imaging. Biosensors, 12(8), 616. 10.3390/bios12080616

Feng, C., Liu, Y., Yuan, Y., Cui, W., Zheng, F., Ma, Y. & Piao, M. (2016). Isoflurane anesthesia exacerbates learning and memory impairment in zinc-deficient APP/PS1 transgenic mice. Neuropharmacology, 111, 119–129. 10.1016/j.neuropharm.2016.08.035

Ferris, C. F. (2022). Applications in Awake Animal Magnetic Resonance Imaging. Frontiers in Neuroscience, 16, 854377. 10.3389/fnins.2022.854377

Ferris, C. F., Smerkers, B., Kulkarni, P., Caffrey, M., Afacan, O., Toddes, S., Stolberg, T. & Febo, M. (2011). Functional magnetic resonance imaging in awake animals. Reviews in the Neurosciences, 22(6), 665–674. 10.1515/rns.2011.050

Fu, Z., Liu, J., Salman, M. S., Sui, J. & Calhoun, V. D. (2023). Functional connectivity uniqueness and variability? Linkages with cognitive and psychiatric problems in children. Nature Mental Health, 1(12), 956–970. 10.1038/s44220-023-00151-8

Gao, Y.-R., Ma, Y., Zhang, Q., Winder, A. T., Liang, Z., Antinori, L., Drew, P. J. & Zhang, N. (2017). Time to wake up: Studying neurovascular coupling and brain-wide circuit function in the un-anesthetized animal. NeuroImage, 153, 382–398. 10.1016/j.neuroimage.2016.11.069

Grandjean, J., Azzinnari, D., Seuwen, A., Sigrist, H., Seifritz, E., Pryce, C. R. & Rudin, M. (2016). Chronic psychosocial stress in mice leads to changes in brain functional connectivity and metabolite levels comparable to human depression. NeuroImage, 142, 544–552. 10.1016/j.neuroimage.2016.08.013

Grandjean, J., Lake, E. M. R., Pagani, M. & Mandino, F. (2024). What N Is N-ough for MRI-Based Animal Neuroimaging? eNeuro, 11(3), ENEURO.0531-23.2024. 10.1523/eneuro.0531-23.2024

Guo, L. Y., Kaustov, L., Brenna, C. T. A., Patel, V., Zhang, C., Choi, S., Halpern, S., Wang, D.-S. & Orser, B. A. (2023). Cognitive deficits after general anaesthesia in animal models: a scoping review. British Journal of Anaesthesia, 130(2), e351–e360. 10.1016/j.bja.2022.10.004

Gutierrez-Barragan, D., Singh, N. A., Alvino, F. G., Coletta, L., Rocchi, F., Guzman, E. D., Galbusera, A., Uboldi, M., Panzeri, S. & Gozzi, A. (2022). Unique spatiotemporal fMRI dynamics in the awake mouse brain. Current Biology. 10.1016/j.cub.2021.12.015

Han, F., Zhao, J. & Zhao, G. (2021). Prolonged Volatile Anesthetic Exposure Exacerbates Cognitive Impairment and Neuropathology in the 5xFAD Mouse Model of Alzheimer’s Disease. Journal of Alzheimer’s Disease, 84(4), 1551–1562. 10.3233/jad-210374

Han, Z., Chen, W., Chen, X., Zhang, K., Tong, C., Zhang, X., Li, C. T. & Liang, Z. (2019). Awake and behaving mouse fMRI during Go/No-Go task. NeuroImage, 188, 733–742. 10.1016/j.neuroimage.2019.01.002

Harris, A. P., Lennen, R. J., Marshall, I., Jansen, M. A., Pernet, C. R., Brydges, N. M., Duguid, I. C. & Holmes, M. C. (2015). Imaging learned fear circuitry in awake mice using fMRI. European Journal of Neuroscience, 42(5), 2125–2134. 10.1111/ejn.12939

He, K., Li, Y., Xiong, W., Xing, Y., Gao, W., Du, Y., Kong, W., Chen, L., Yang, X. & Dai, Z. (2024). Sevoflurane exposure accelerates the onset of cognitive impairment via promoting p-Drp1S616-mediated mitochondrial fission in a mouse model of Alzheimer’s disease. Free Radical Biology and Medicine, 225, 699–710. 10.1016/j.freeradbiomed.2024.10.301

Hori, Y., Schaeffer, D. J., Gilbert, K. M., Hayrynen, L. K., Cléry, J. C., Gati, J. S., Menon, R. S. & Everling, S. (2020). Altered Resting-State Functional Connectivity Between Awake and Isoflurane Anesthetized Marmosets. Cerebral Cortex, 30(11), 5943–5959. 10.1093/cercor/bhaa168

Kim, B., Boes, J. L., Bland, P. H., Chenevert, T. L. & Meyer, C. R. (1999). Motion correction in fMRI via registration of individual slices into an anatomical volume. Magnetic Resonance in Medicine, 41(5), 964–972. 10.1002/(sici)1522-2594(199905)41:5&964::aid-mrm16>3.0.co;2-d

Kroll, T., Kornadt-Beck, N., Oskamp, A., Elmenhorst, D., Touma, C., Palme, R. & Bauer, A. (2021). Additional Assessment of Fecal Corticosterone Metabolites Improves Visual Rating in the Evaluation of Stress Responses of Laboratory Rats. Animals, 11(3), 710. 10.3390/ani11030710

Lindhardt, T. B., Gutiérrez-Jiménez, E., Liang, Z. & Hansen, B. (2022). Male and Female C57BL/6 Mice Respond Differently to Awake Magnetic Resonance Imaging Habituation. Frontiers in Neuroscience, 16, 853527. 10.3389/fnins.2022.853527

Low, L. A., Bauer, L. C., Pitcher, M. H. & Bushnell, M. C. (2016). Restraint training for awake functional brain scanning of rodents can cause long-lasting changes in pain and stress responses. PAIN, 157(8), 1761–1772. 10.1097/j.pain.0000000000000579

Luppi, A. I., Golkowski, D., Ranft, A., Ilg, R., Jordan, D., Bzdok, D., Owen, A. M., Naci, L., Stamatakis, E. A., Amico, E. & Misic, B. (2025). General anaesthesia decreases the uniqueness of brain functional connectivity across individuals and species. Nature Human Behaviour, 9(5), 987– 1004. 10.1038/s41562-025-02121-9

Madularu, D., Mathieu, A. P., Kumaragamage, C., Reynolds, L. M., Near, J., Flores, C. & Rajah, M. N. (2017). A non-invasive restraining system for awake mouse imaging. Journal of Neuroscience Methods, 287, 53–57. 10.1016/j.jneumeth.2017.06.008

Mandino, F., Cerri, D. H., Garin, C. M., Straathof, M., Tilborg, G. A. F. van, Chakravarty, M. M., Dhenain, M., Dijkhuizen, R. M., Gozzi, A., Hess, A., Keilholz, S. D., Lerch, J. P., Shih, Y.-Y. I. & Grandjean, J. (2020). Animal Functional Magnetic Resonance Imaging: Trends and Path Toward Standardization. Frontiers in Neuroinformatics, 13, 78. 10.3389/fninf.2019.00078

Mandino, F., Vrooman, R. M., Foo, H. E., Yeow, L. Y., Bolton, T. A. W., Salvan, P., Teoh, C. L., Lee, C. Y., Beauchamp, A., Luo, S., Bi, R., Zhang, J., Lim, G. H. T., Low, N., Sallet, J., Gigg, J., Lerch, J. P., Mars, R. B., Olivo, M., … Grandjean, J. (2021). A triple-network organization for the mouse brain. Molecular Psychiatry, 1–8. 10.1038/s41380-021-01298-5

Mandino, F., Vujic, S., Grandjean, J. & Lake, E. M. R. (2023). Where do we stand on fMRI in awake mice? *Cerebral Cortex*, bhad478. 10.1093/cercor/bhad478

Mardini, F., Tang, J. X., Li, J. C., Arroliga, M. J., Eckenhoff, R. G. & Eckenhoff, M. F. (2017). Effects of propofol and surgery on neuropathology and cognition in the 3xTgAD Alzheimer transgenic mouse model. British Journal of Anaesthesia, 119(3), 472–480. 10.1093/bja/aew397

Maron-Katz, A., Vaisvaser, S., Lin, T., Hendler, T. & Shamir, R. (2016). A large-scale perspective on stress-induced alterations in resting-state networks. Scientific Reports, 6(1), 21503. 10.1038/srep21503

Miao, H., Dong, Y., Zhang, Y., Zheng, H., Shen, Y., Crosby, G., Culley, D. J., Marcantonio, E. R. & Xie, Z. (2018). Anesthetic Isoflurane or Desflurane Plus Surgery Differently Affects Cognitive Function in Alzheimer’s Disease Transgenic Mice. Molecular Neurobiology, 55(7), 5623–5638. 10.1007/s12035-017-0787-9

Power, J. D., Barnes, K. A., Snyder, A. Z., Schlaggar, B. L. & Petersen, S. E. (2012). Spurious but systematic correlations in functional connectivity MRI networks arise from subject motion. NeuroImage, 59(3), 2142–2154. 10.1016/j.neuroimage.2011.10.018

Power, J. D., Plitt, M., Gotts, S. J., Kundu, P., Voon, V., Bandettini, P. A. & Martin, A. (2018). Ridding fMRI data of motion-related influences: Removal of signals with distinct spatial and physical bases in multiecho data. Proceedings of the National Academy of Sciences, 115(9), E2105– E2114. 10.1073/pnas.1720985115

Rowland, N. E. & Toth, L. A. (2019). Analytic and Interpretational Pitfalls to Measuring Fecal Corticosterone Metabolites in Laboratory Rats and Mice. Comparative Medicine, 69(5), 337–349. 10.30802/aalas-cm-18-000119

Russo, G., Helluy, X., Behroozi, M. & Manahan-Vaughan, D. (2021). Gradual Restraint Habituation for Awake Functional Magnetic Resonance Imaging Combined With a Sparse Imaging Paradigm Reduces Motion Artifacts and Stress Levels in Rodents. Frontiers in Neuroscience, 15, 805679. 10.3389/fnins.2021.805679

Schuetze, S., Manig, A., Ribes, S. & Nau, R. (2019). Aged mice show an increased mortality after anesthesia with a standard dose of ketamine/xylazine. Laboratory Animal Research, 35(1), 8. 10.1186/s42826-019-0008-y

Smith, J., Wilkey, E., Clarke, B., Shanley, L., Men, V., Fair, D. & Sabb, F. W. (2022). Can this data be saved? Techniques for high motion in resting state scans of first grade children. Developmental Cognitive Neuroscience, 58, 101178. 10.1016/j.dcn.2022.101178

Soares, J. M., Magalhães, R., Moreira, P. S., Sousa, A., Ganz, E., Sampaio, A., Alves, V., Marques, P. & Sousa, N. (2016). A Hitchhiker’s Guide to Functional Magnetic Resonance Imaging. Frontiers in Neuroscience, 10, 515. 10.3389/fnins.2016.00515

Szilagyi, K., Zieger, M. A., Li, J. & Kacena, M. A. (2018). Improving Post-Operative Outcomes in Aged and Diabetic Obese Mice. Laboratory Animal Science Professional, 6(3), 65–67.

Takata, N., Sugiura, Y., Yoshida, K., Koizumi, M., Hiroshi, N., Honda, K., Yano, R., Komaki, Y., Matsui, K., Suematsu, M., Mimura, M., Okano, H. & Tanaka, K. F. (2018). Optogenetic astrocyte activation evokes BOLD fMRI response with oxygen consumption without neuronal activity modulation. Glia, 66(9), 2013–2023. 10.1002/glia.23454

Tang, J. X. & Eckenhoff, M. F. (2013). Anesthetic effects in Alzheimer transgenic mouse models. Progress in Neuro-Psychopharmacology and Biological Psychiatry, 47, 167–171. 10.1016/j.pnpbp.2012.06.007

Tang, J. X., Mardini, F., Caltagarone, B. M., Garrity, S. T., Li, R. Q., Bianchi, S. L., Gomes, O., Laferla, F. M., Eckenhoff, R. G. & Eckenhoff, M. F. (2011). Anesthesia in presymptomatic Alzheimer’s disease: A study using the triple-transgenic mouse model. Alzheimer’s & Dementia, 7(5), 521–531.e1. 10.1016/j.jalz.2010.10.003

Tsurugizawa, T. & Yoshimaru, D. (2021). Impact of anesthesia on static and dynamic functional connectivity in mice. NeuroImage, 241, 118413. 10.1016/j.neuroimage.2021.118413

Wang, S., Peterson, D. J., Gatenby, J. C., Li, W., Grabowski, T. J. & Madhyastha, T. M. (2017). Evaluation of Field Map and Nonlinear Registration Methods for Correction of Susceptibility Artifacts in Diffusion MRI. Frontiers in Neuroinformatics, 11, 17. 10.3389/fninf.2017.00017

Winkler, A. M., Ridgway, G. R., Webster, M. A., Smith, S. M. & Nichols, T. E. (2014). Permutation inference for the general linear model. NeuroImage, 92(100), 381–397. 10.1016/j.neuroimage.2014.01.060

Wu, T., Wang, X., Zhang, R., Jiao, Y., Yu, W., Su, D., Zhao, Y. & Tian, J. (2020). Mice with pre-existing tumors are vulnerable to postoperative cognitive dysfunction. Brain Research, 1732, 146650. 10.1016/j.brainres.2020.146650

Xu, N., LaGrow, T. J., Anumba, N., Lee, A., Zhang, X., Yousefi, B., Bassil, Y., Clavijo, G. P., Sharghi, V. K., Maltbie, E., Meyer-Baese, L., Nezafati, M., Pan, W.-J. & Keilholz, S. (2022). Functional Connectivity of the Brain Across Rodents and Humans. Frontiers in Neuroscience, 16, 816331. 10.3389/fnins.2022.816331

Xu, W., Pei, M., Zhang, K., Tong, C., Bo, B., Feng, J., Zhang, X.-Y. & Liang, Z. (2022). A systematically optimized awake mouse fMRI paradigm. bioRxiv, 2022.11.16.516376. 10.1101/2022.11.16.516376

Yun, Y. L. & Grandjean, J. (2020). Mouse_rest_awake. 10.18112/openneuro.ds001653.v1.0.2

Zeng, H., Jiang, Y., Beer-Hammer, S. & Yu, X. (2022). Awake Mouse fMRI and Pupillary Recordings in the Ultra-High Magnetic Field. Frontiers in Neuroscience, 16, 886709. 10.3389/fnins.2022.886709

Zhang, C., Zhang, Y., Shen, Y., Zhao, G., Xie, Z. & Dong, Y. (2017). Anesthesia/Surgery Induces Cognitive Impairment in Female Alzheimer’s Disease Transgenic Mice. Journal of Alzheimer’s Disease, 57(2), 505–518. 10.3233/jad-161268

Zhao, W., Makowski, C., Hagler, D. J., Garavan, H. P., Thompson, W. K., Greene, D. J., Jernigan, T. L. & Dale, A. M. (2023). Task fMRI paradigms may capture more behaviorally relevant information than resting-state functional connectivity. NeuroImage, 270, 119946. 10.1016/j.neuroimage.2023.119946

Zhou, S., Cui, X., Chen, J., Luo, M., Ouyang, W., Tong, J., Xie, Z. & Le, Y. (2024). Single exposure to anesthesia/surgery in neonatal mice induces cognitive impairment in young adult mice. Free Radical Biology and Medicine, 214, 184–192. 10.1016/j.freeradbiomed.2024.02.017

