## Supplementary Figures and Tables for "A non-invasive approach to awake mouse fMRI compatible with multi-modal techniques"

Supplementary Table 1. RABIES (version 0.5.1) processing stage commands.

| Processing Stage | Parameters |
|---|---|
| Preprocessing | --anat_robust_inho_cor apply=true,masking=false,brain_extraction=false,template_registration=SyN<br>--commonspace_resampling 0.2x0.2x0.2<br>--bold2anat_coreg masking=true,brain_extraction=false,registration=SyN |
| Confound Correction | --smoothing_filter 0.3<br>--conf_list mot_6<br>--frame_censoring FD_censoring=true,FD_threshold=0.15,DVARS_censoring=false,minimum_timepoint=200 |
| Analysis | --data_diagnosis |

Supplementary Table 2. Percentage of volumes censored for each group using motion thresholds as outlined in Supplementary Table 1. The first 4 rows represent the data divided based on acclimation group. The last 5 rows represent the non-invasive data grouped by average framewise displacement (FD) per run. The range of average run FD is reported under each group (parentheses indicate exclusivity, square brackets indicate inclusivity). The last column, Average Volumes Censored Per Subject, indicates the number of censored volumes (average ± standard deviation) for each subject in the different acclimation groups. The non-invasive groups were not significantly different from one another (1-way ANOVA, $p > 0.05$). The last column was not computed for the motion groups (last 5 rows) since the same subject may appear in different groups depending on the motion level for that run.

| Group | Total Runs | Censored Volumes | Runs Excluded | Average Volumes Censored Per Subject |
|---|---|---|---|---|
| Group 1: 4-Day (0.039 – 0.13 mm) | 40 | 22.3% | 4 (10.0%) | 25.1% ± 16.2% |
| Group 2: 9-Day (0.015 – 0.14 mm) | 80 | 13.2% | 4 (5.0%) | 14.1% ± 6.3% |
| Group 3: 13-Day (0.014 – 0.16 mm) | 102 | 17.2% | 10 (9.8%) | 16.2% ± 8.4% |
| Group 4: Headpost (0.018 – 0.027 mm) | 14 | 0.0% | 0 (0.0%) | 0% ± 0% |
| Motion Level 1 [0.000 – 0.030 mm) | 26 | 0.3% | 0 (0.0%) | |
| Motion Level 2 [0.030 – 0.050 mm) | 59 | 4.5% | 0 (0.0%) | |
| Motion Level 3 [0.050 – 0.070 mm) | 53 | 12.1% | 0 (0.0%) | |
| Motion Level 4 [0.070 – 0.090 mm) | 44 | 23.4% | 0 (0.0%) | |
| Motion Level 5 [0.090 – 0.110 mm) | 40 | 43.9% | 18 (45.0%) | |

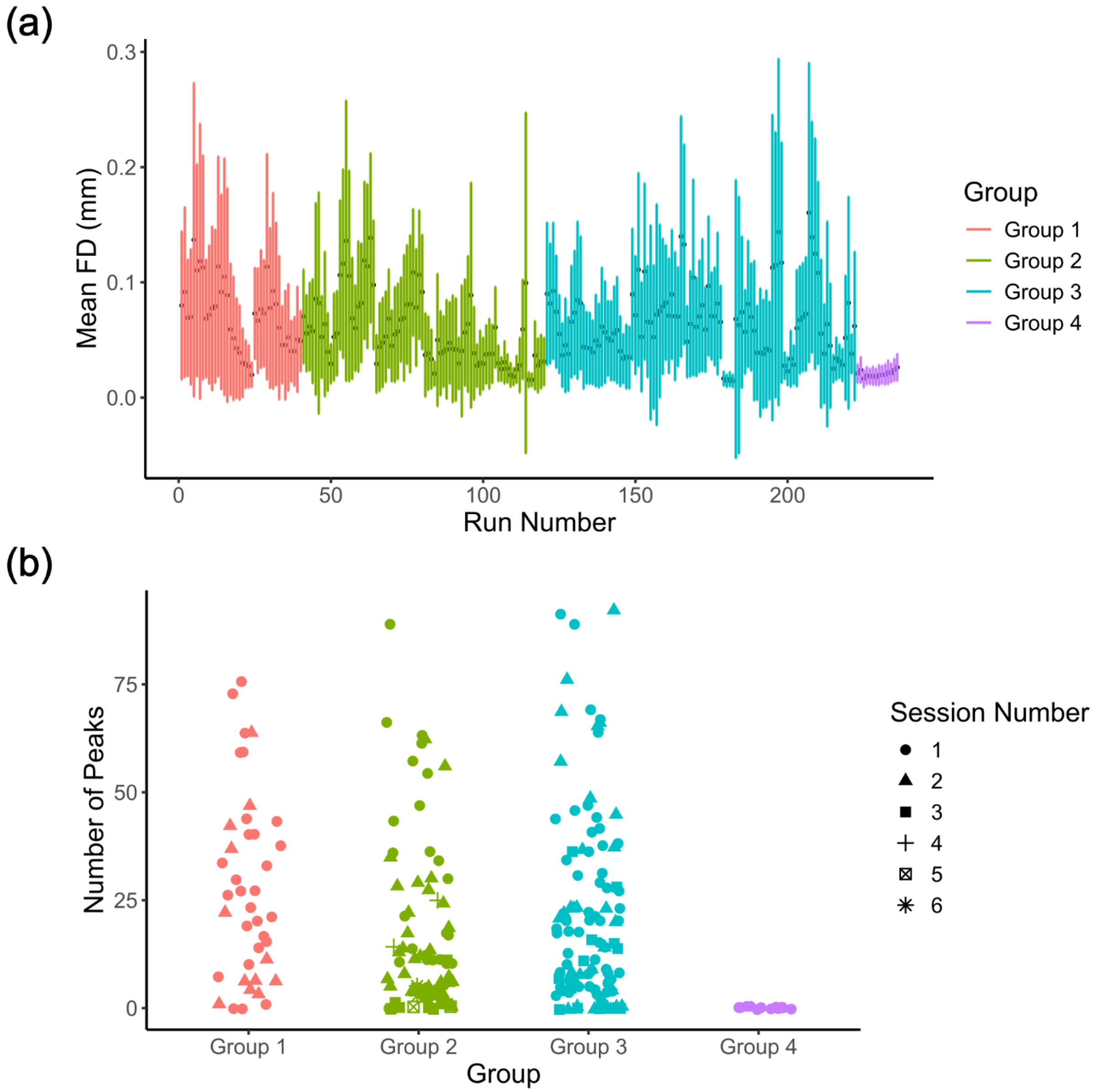


Supplementary Figure 1. (a) Mean run FD as a function of run number. Runs are split up based on acclimation protocol (Group1 = 4-day, Group 2 = 9-day, Group 3 = 13-day, and Group 4 = headpost) and further sorted by increasing session number. Dots represent the average run FD and error bars represent the standard deviation within the run. (b) Number of peaks of excessive motion (i.e., above FD threshold of 0.15 mm) for each run in the 4 groups. Marker shape is used to differentiate the session number as indicated by the legend to the right.

Supplementary Table 3. Number and percentage of volumes censored of all data analyzed using different censoring thresholds.

| Censoring Threshold | Volumes Remaining |
| --- | --- |
| No threshold | 97,200 (100.0%) |
| 150 μm | 78,795 (81.1%) |
| 100 μm | 62,562 (64.4%) |
| 50 μm | 33,224 (34.2%) |

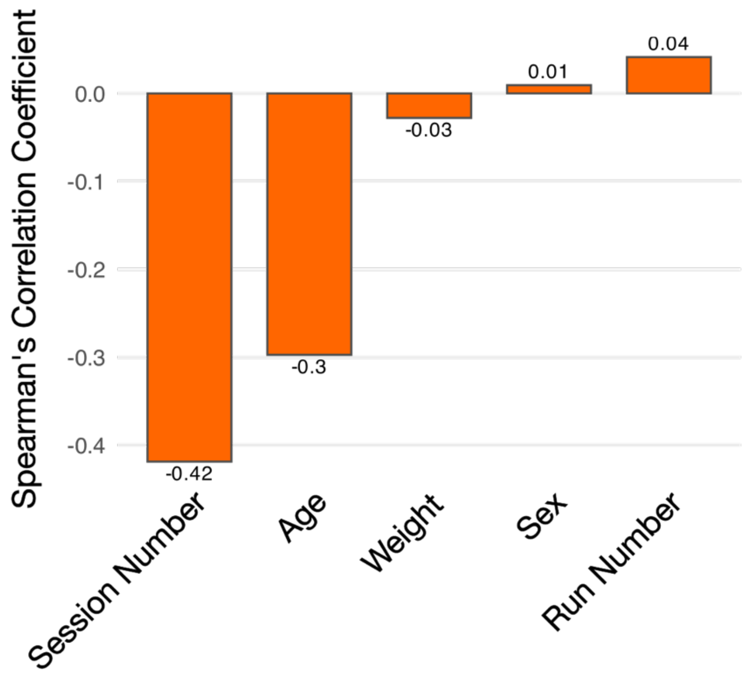


Supplementary Figure 2. Spearman's rank correlation coefficient calculated between average run FD value and various metrics at the time of the acquisition including session number, age, weight, sex and run number. Session number is the number of previous scanning sessions the mouse has undergone (i.e., potential measure of acclimation), age and weight are at the time of the corresponding run acquisition, and the run number is the number of runs the mouse has undergone in the current scanning session.

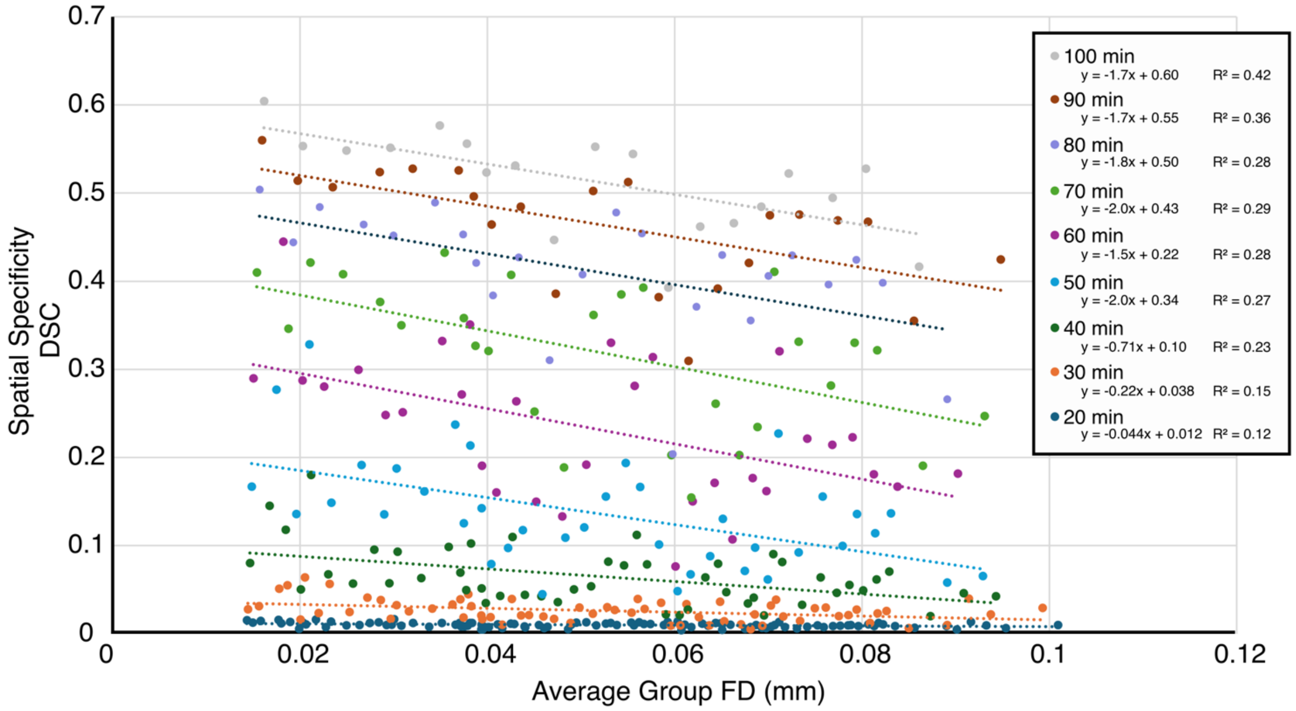


Supplementary Figure 3. Relationship between spatial specificity (Dice Similarity Coefficient, DSC) and average group framewise displacement (FD) for different group cumulative acquisition times. All functional runs were aggregated and sorted based on the average FD value. Group network spatial specificity was determined for all groups against the prior somatomotor network, and the results were plotted. Linear trendlines were fitted to each cumulative acquisition time groupings. The trendline equations and coefficients of determination ($R^2$) are displayed in the legend under each grouping.

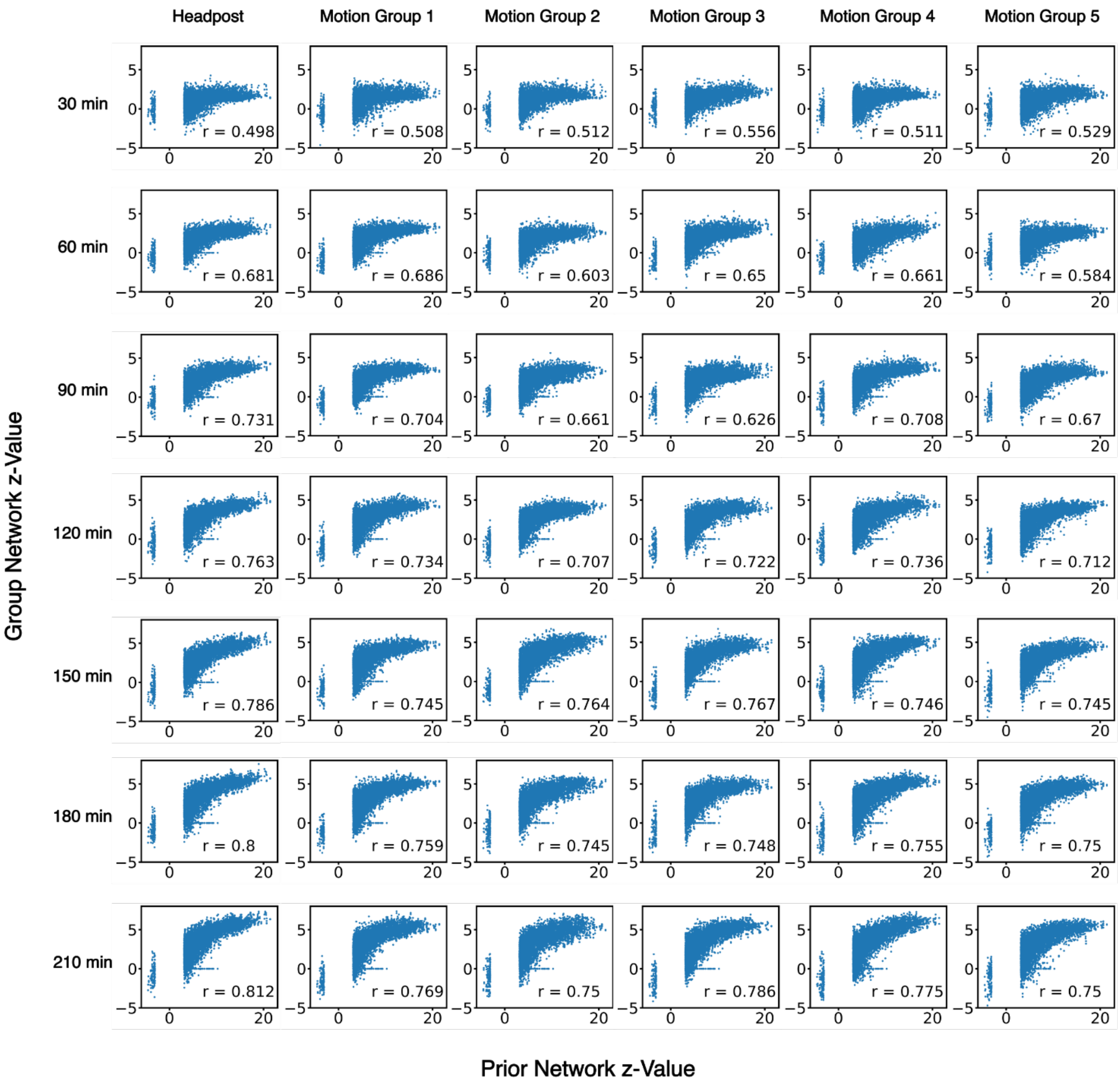


Supplementary Figure 4. Correspondence between z-values of the prior somatomotor network and corresponding group networks on a voxel-wise basis for voxels in the prior network region. One sample from each group network is shown. Each column represents a different motion level and each row represents a different cumulative group acquisition time. The gap between -3.1 and 3.1 results from masking the prior network for voxels with values >= 3.1 and <= -3.1 and selecting those voxels for the correlation analysis. The corresponding Pearson Correlation coefficient (r value) is annotated on each plot.

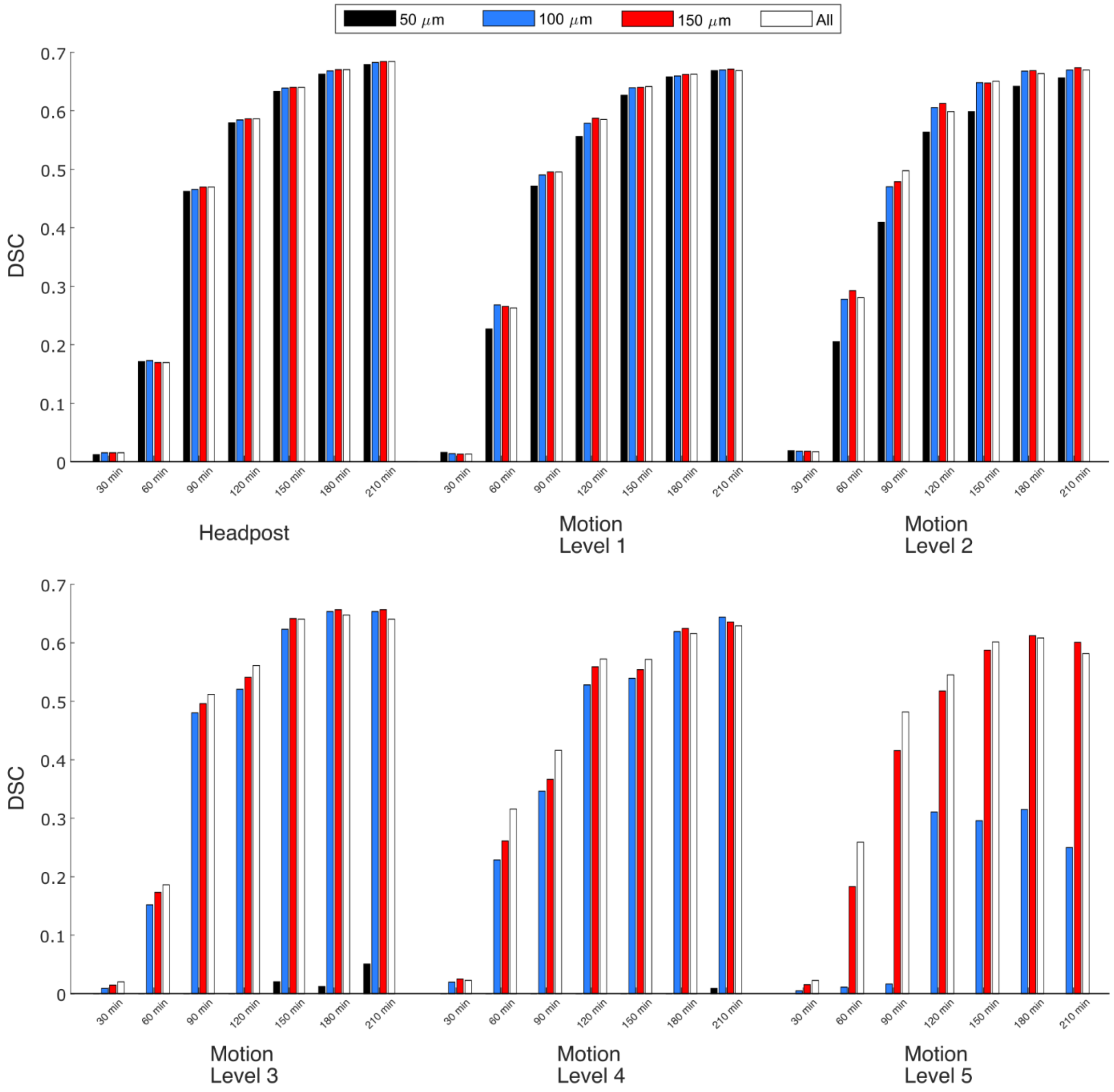


Supplementary Figure 5. Effect of censoring threshold on network quality. The Dice Similarity Coefficient (DSC) was used to measure network overlap with the prior somatomotor network. The categories on the x-axis represent the cumulative group acquisition time used to create each group network for the headpost group and the non-invasive groups at different motion levels (i.e., Motion Level 1-5). The three censoring thresholds used were 50 μm in black, 100 μm in blue, 150 μm in red, and no threshold (i.e., “All”) in white.

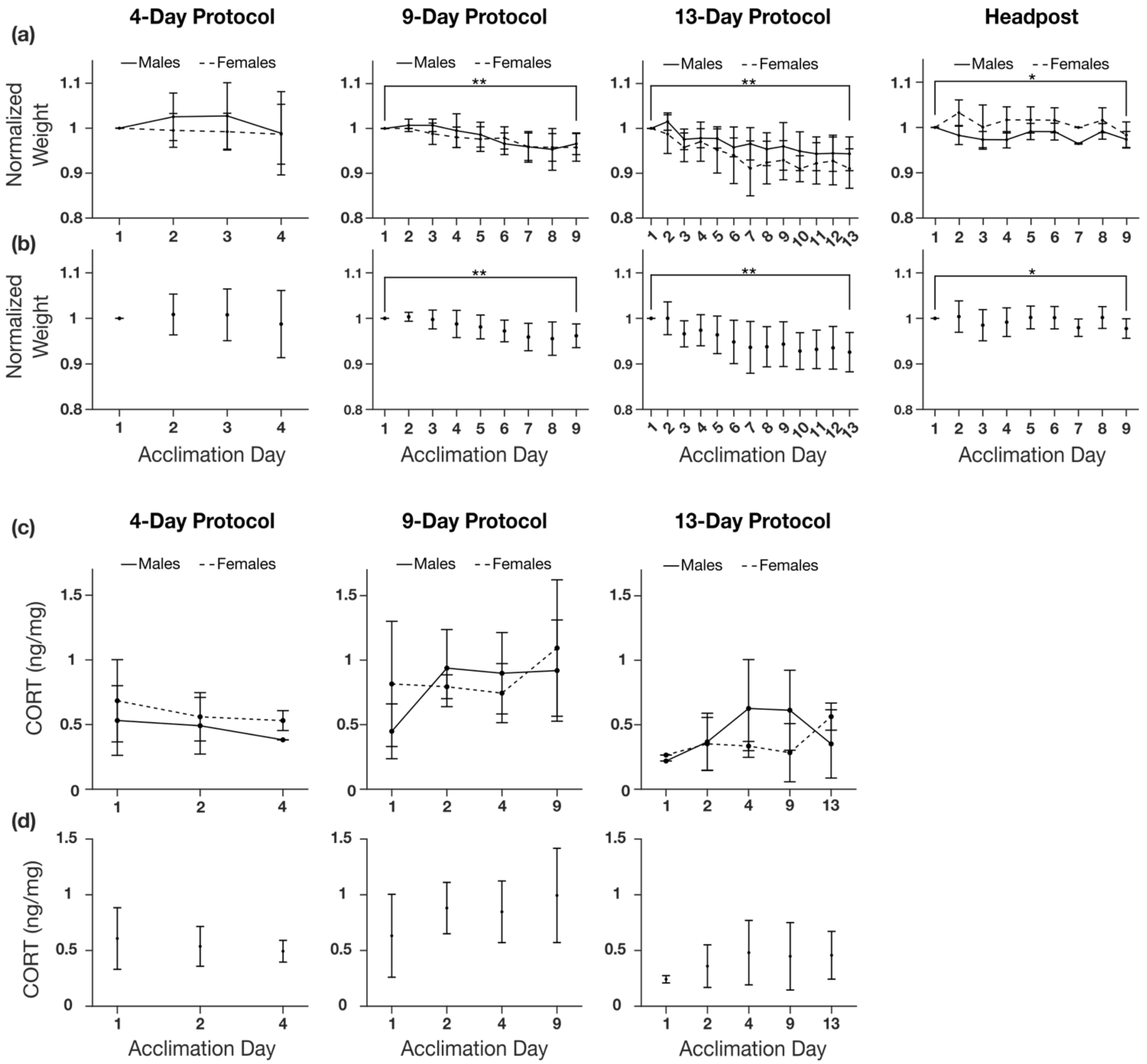


Supplementary Figure 6. Weight and fecal corticosterone (CORT) metabolite measured over 3 acclimation protocols. (a) Normalized weight split up by sex for the 4-day, 9-day, 13-day and headpost groups. The weight on the last day was significantly less than the first day for the 9-day, 13-day and headpost (only males) but not for the 4-day protocol (paired t-test, $p < 0.05$). (b) Same as (a) except both sexes aggregated. (c) CORT split up by sex for the 4-day, 9-day, and 13-day groups. (d) Same as (c) except both sexes aggregated. CORT concentrations are displayed in ng of CORT per mg of fecal matter. There were no significant CORT changes across acclimation sessions (one-way ANOVA, $p > 0.05$). Data are displayed as group means (data points) and standard deviation (error bars).

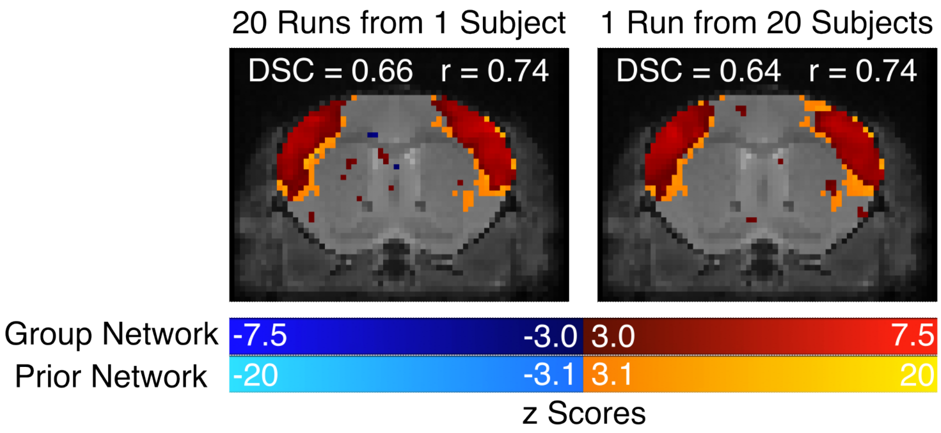


Supplementary Figure 7. Sample slice from a somatomotor network group map comprised of 20 runs from the same subject (left) and another group comprised of 20 runs from different subjects (1 run from 20 different subjects) (right). Group networks (dark red/dark blue) are overlayed on top of the prior somatomotor network (orange/light blue).

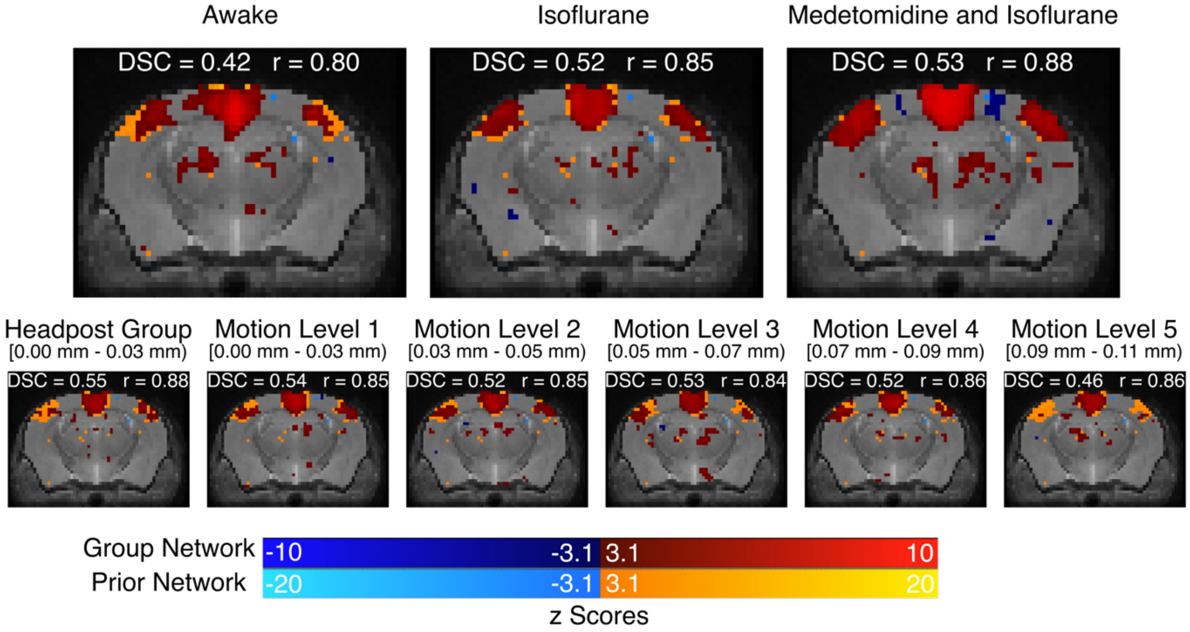


Supplementary Figure 8. Comparison of group DMN maps from an external dataset compared with the data from this manuscript with the same cumulative acquisition time (180 min). The top row shows the maps from the external dataset in subjects scanned awake (left), using isoflurane (centre), and using both medetomidine and isoflurane (right). The DSC and amplitude correlation values are annotated above each map.
